# The observation of birds as informative tools to develop citizen science: Contributions of a 10-year record in the state of Chihuahua, Mexico

**DOI:** 10.1101/704676

**Authors:** Fidel González-Quiñones, Luis Roberto Granados, José Manuel Jurado Ruiz, Javier Tarango, Juan D. Machin-Mastromatteo, José Refugio Romo-González

## Abstract

This article analyses historical data from observations made to birds in breeding, throughout two routes with urban characteristics and during a consecutive period of 10 years (2009-2018), following a precise methodology designed by the North American Breeding Bird Survey. The analyzed routes are officially registered in the Mexican Commission for Biodiversity’s Knowledge and Use, the United States Geological Survey Patuxent Wildlife Research Center, and the Canadian Wildlife Service Research Centre. The observations were made by citizens without formal professional education; hence the results may be considered within the framework of citizen science. Their contributions provided important data for decision-making regarding environmental issues, since the presence of birds is considered one of the main indicators on health conditions of an ecosystem. Data analysis identified two basic conditions: (i) a reduction of the 23% in the number of species found, as many of them disappeared during counting; and (ii) the significant increase in population of other species, three of pigeons among them. Apart from the study of bird behavior in the routes with urban characteristics, the article acknowledges the lack of connection and use of the information produced from monitoring for decision-making and education regarding environmental issues. Therefore, we consider crucial to create scientific observatories, both available to experts in the field and to the general population, as the ultimate purpose would be the production of citizen science.

## 1. Introduction

Birdwatching is an activity where a person or a group of people on their own get organized to identify wild or feral birds that live in a specific environment. The main characteristics for the process of birdwatching are: (i) it can be performed by amateurs or experts; (ii) in general, it does not require specialized or scientific training; and (iii) it can be initiated with a relatively small financial investment.

In order to correctly conduct this type of process, bird watchers make sure to interact the least possible with the fauna and the sighting places, restricting themselves to exclusively register their observations with photographs, videos, audios and notes, and they may also use simple manuals and guides [1]. While watchers develop greater experience, they might need some tools that allow them to conduct observations in a more systematic way and get more evidence from them. Among the main tools used, there are binoculars, cameras that include lenses of 300 millimeters at least, and specialized guides [2].

For the case of this investigation, the study was conducted in the city of Chihuahua, capital of the state of Chihuahua to the north of Mexico, neighboring with the United States. In the local environment, the most experienced bird watchers carry out the functions of training and leading amateur observers, and they perform several activities regarding education and training, such as courses for the identification of birds, collective photographical expositions, and training. Also, those who have experience in falconry, rehabilitate birds of prey at the request of the Federal Government of Mexico, through the Federal Government’s Environmental Protection Agency [3]. In addition to the members of this investigation, there is a well-known group in the city of Chihuahua: ‘Colectivo de Aves’ (Bird Collective), which conducts different cultural and ecological activities [4], and some members of this group have been trained and certified in monitoring and birdwatching by the Mexican Commission for Biodiversity’s Knowledge and Use (CONABIO) [5], the United States Geological Survey Patuxent Wildlife Research Center [6] and the Canadian Wildlife Service Research [7].

Although there are limited supporting options for developing activities related to environmental issues in Mexico, it is important to point out that many of these activities emerge from voluntary initiatives. In the particular case of those participating in the observation and supervision of birds, they conduct these activities because of the esthetic satisfaction they find in them and because of a personal interest, given the interaction with the birds, the environment and other participants. Beyond this, there are some strategies for the conservation and usage of the species observed, as well as for the knowledge of the environmental impact of the anthropogenic activities in the local, regional and global level [8].

### 1.1. Urban infrastructure and bird observation

In the context of the education for the sustainability of life, birdwatching activities require the participation of people who learn and get involved with topics related to the avian life that coexist in their contexts, which leads to promote several elements to improve the environment, such as encouraging the conservation of species and improving cities accordingly, as they should also be suitable habitats for other living beings [9]. These remarks on urban infrastructure and human activities also include other points of view related to humans and their relationship with birds, such as the habit of feeding them and giving them shelter [10]; and providing citizens with tips for their diet and care [11].

It is not necessary to be an expert in the natural sciences to observe birds. Many of the people who do it are retired, kids or people whose professional life is not connected with this field, such as doctors, architects, chemists, engineers, entrepreneurs and geographers [12]. However, some people involved in these groups have enough technical training and knowledge to perform such activities, which are based on a scientific methology for monitoring regular routes, annually and periodically, to report the influence of birds in a specific location [13]. An example of this is the Sitio Ramsar Vado de Meoqui, a town located in the state of Chihuahua, which is considered a protected area under an international environmental agreement that enforces the conservation and rational use of the wetlands through local, regional and national actions for contributing to sustainable development in the world [14]. These are transcendental remarks, since the mentioned town is located within an urban area, which has a record of serious damages in its ecology, due to inappropriate urban planning and human activities that produce waste and contamination [15].

In the context of a primarily urban modern society, 74.2% of the Mexican population lives in cities of more than 15 thousand inhabitants [16]. Under such conditions, it is compulsory to raise awareness and to re-educate people regarding sustainable development, by carrying out activities such as the systematic and participative observation of the ecological dynamics in populations such as birds.

Within the actions based on systematic methodologies for birdwatching, it is expected that observers who start these activities as a pastime, to eventually perform them in a more constant and formal way. An example of this is quoted in the report that documents the observation of birds as a serious proposal for the environmental education to teachers and students in the protected natural areas of the Parque Natural Municipal de Saltos Küpper and the Reserva Privada de Vida Silvestre Virgen de Paticuá of the town Eldorado, Colombia, where bird data collection were carried out with the support of common citizens [17]. This example is important because Colombia has the largest diversity of birds in the world, and it has a great ornithological tradition, although birdwatching as a pastime is a recent activity [18].

Guayaquil city (Ecuador) has another experience with a different approach, but related to birdwatching, education, ethics and cultural coexistence in urban areas. In this city, 130 species of birds were identified in a study conducted during 20 months, which also contributed to the building of pathways, tourist facilities, rest areas, viewpoints, piers, as well as accessible infrastructure, lightning and parking lots [19].

In the literature review, we identified several birdwatching projects in urban and semi urban areas under different modalities and dimensions; such as those associating ethics with birds, peoples and cultures, as well as the observation and reflection upon birds’ ecological and cultural aspects [20]. Other studies were conducted on a single species, by analyzing distribution models that employ satellite tracking software and observation through computing platforms [21].

The study and tracking of birds in urban and semi urban environments have become more popular activities for different reasons, such as the closeness of the observation sites to the watchers’ homes [22], the opportunity to become familiar with the species of the area and identifying their main characteristics [23]; and learning about the urban features and their influence in birds’ behavior, for instance in desert environments [24].

Sometimes, birdwatching is conducted in semi urban areas, which are under some sort of special surveillance, such as government installations or inside educational institutions. The experiences identified in Latin America are: (i) inside the college campus ‘El Cerillo’ in Universidad Autónoma del Estado de México, which reported 141 species [25]; (ii) at Universidad del Amazonas in Colombia, which reported 62 species [26]; (iii) at Universidad de Asunción (Paraguay), with 77 species [27]; (iv) at Universidad de El Salvador, with 64 species [28]; and (v) 36 species reported in Centro Regional Universitario de Colón (Panama) [29]. Some of these cases have studied the collisions of birds into windows, which is considered an important aspect, because a great number of birds die for this reason [30].

Another important feature of birdwatching is their bond with the tourist activity, for instance, this type of activity in Cuba has gone from an initial extractive approach to a relatively innocuous activity for birds, and it has turned into an important economic activity; however, this country does not have a record of the impact caused by tourism to birdwatching [31]. In the state of Oaxaca (Mexico), birdwatching is an ecotouristim activity offered by people or companies to tourists mainly from the United States and Canada [32]. As a result, the development of formal training programs for touristic guides has been proposed, for making such activities environmentally sustainable [33].

In Mexico, there are governmental institutions that work in a methodic way in the observation of birds, such as the Commission for Biodiversity’s Knowledge and Use (CONABIO) [5], which fosters birdwatching and the funding for the people involved in it. The data gathered in the monitoring activities supported by CONABIO allow identifying the density of birds’ population and health indicators of the ecosystems, such as the rise or fall on the number of individuals for a specific species, and also detecting environmental alteration elements, such as droughts or groundwater depletion, harmful fertilizers, and the commerce or illegal trade of species [34]. In Mexico, birdwatching was recently included in a legal environmental study related to the constructions of the new airport in Mexico City, as the presence of 131 species was reported in the construction area, which might have an influence in plane crashes [35].

In the particular case of the state of Chihuahua, the scenario of the present study, we identified the following studies: (i) in the Ramsar zone known as ‘Vado de Meoqui’ in the riverbed of Río San Pedro [36]; (ii) the identification of 30 species of birds through a study of variability about the frequency of sighting of seabirds, in relation to the months of year [13]; (iii) the observation of species of little frequency through 36 points of observation [37]; and (iv) the observation of birds through the presence of nests and hatchings of the American avocet (Recurvirostra americana), which was considered a species present in the area as a result of migration and not permanent, e.g. not born in Chihuahua [38].

## 2. Materials and Methods

This research was based on birdwatching data gathered during 10 continuous years (2009-2018) in two semi urban routes of observation located in the state of Chihuahua: La Regina (code 058) and Santa Mónica (code 059). These routes are registered by CONABIO, the United States Geological Survey Patuxent Wildlife Research Center, and the Canadian Wildlife Service Research.

Each route is 40 kilometers long and they consist of 50 monitoring stops every 800 meters. Both routes pass through one urban area and another semi urban area, thus they are habitats that are directly influenced by human activities, such as dwellings, agricultural and familiar vehicles, parks, gardens, croplands, and irrigation ditches. La Regina is the most affected by land use change. Figure 1 shows the routes in the map with their 50 monitoring stops, which were mapped by using their coordinates.

**Fig 1.**
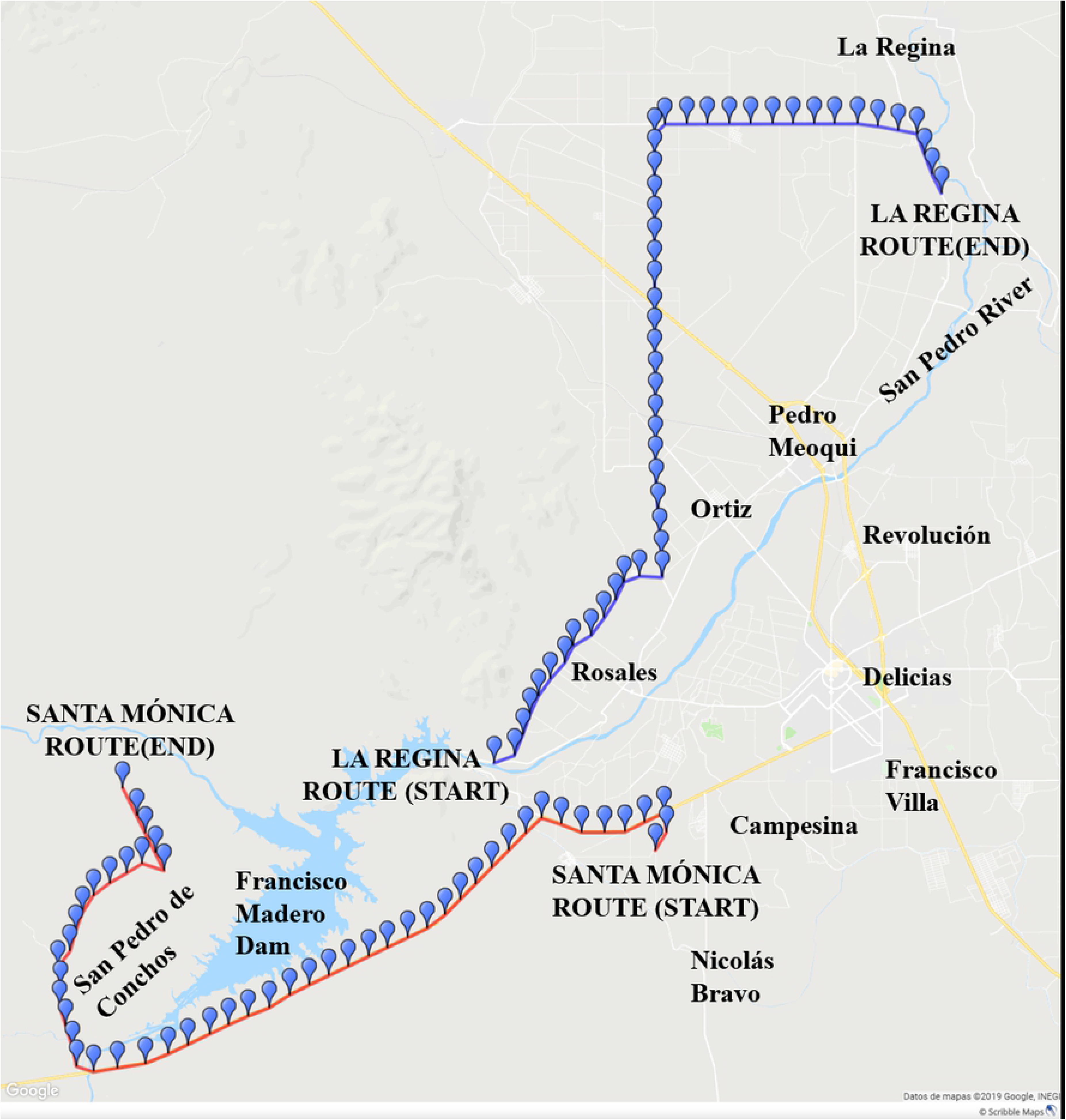
La Regina and Santa Mónica routes in the map

The beginning of each record in the routes was established between 5:00 and 5:30 am, and the monitoring points were marked with a physical description and their GPS coordinates. Also, it is common to place signals and visible marks for an easy identification of the stops every year. Before, during, and at the end of each tracking, the climatological conditions (temperature, wind speed, and cloudiness) are recorded. The monitoring was made every year within the period of May-July, when the breeding season starts and migratory species have already travelled to their nesting places.

The protocol for marking the routes consisted of establishing 50 points of observation separated from each other every 800 meters and geographically located by their physical description and GPS coordinates. At each point, a chronometer was used to measure a period of three minutes and an approximate radius of 400 meters was monitored to record all the bird species observed and heard. This protocol was repeated annually, starting in the same point and at the same hour, to maintain uniformity in sampling conditions.

## 3. Analysis of the results

The descriptive analysis of the species found was made from global data of both routes observed, with its corresponding correlations; using tables and graphics by route, which show the increase and decrease of species, as well as an analysis by number of species. During the 10-year period, a total of 139 species were identified and observed in both routes, Table 1 compiles species common names in English, their scientific names, and the longitudinal behavior from 2009 to 2018, with the corresponding total of sightings per species.

**Table 1.** Total descriptive analysis of the species found.

| No. | Common names (English) | Scientific names | Years of observation |  |  |  |  |  |  |  |  |  |  |
| --- | --- | --- | --- | --- | --- | --- | --- | --- | --- | --- | --- | --- | --- |
|  |  |  | 2009 | 2010 | 2011 | 2012 | 2013 | 2014 | 2015 | 2016 | 2017 | 2018 | TOTAL |
| 1 | *White-winged Dove* | *Zenaida asiática* | 462 | 832 | 830 | 987 | 843 | 897 | 1527 | 1315 | 1518 | 1383 | 10594 |
| 2 | *Mourning Dove* | *Zenaida macroura* | 664 | 729 | 910 | 980 | 915 | 729 | 957 | 1260 | 1555 | 1371 | 10070 |
| 3 | *Red-winged Blackbird* | *Agelaius phoeniceus* | 326 | 439 | 432 | 450 | 496 | 456 | 666 | 552 | 618 | 697 | 5132 |
| 4 | *Great-tailed Grackle* | *Quiscalus mexicanus* | 589 | 603 | 320 | 410 | 342 | 412 | 209 | 362 | 594 | 479 | 4320 |
| 5 | *House Sparrow* | *Passer domesticus* | 264 | 406 | 230 | 187 | 193 | 279 | 417 | 462 | 401 | 455 | 3294 |
| 6 | *Inca Dove* | *Columbina inca* | 131 | 73 | 92 | 100 | 62 | 128 | 179 | 358 | 810 | 649 | 2582 |
|  |  |  | 2009 | 2010 | 2011 | 2012 | 2013 | 2014 | 2015 | 2016 | 2017 | 2018 | TOTAL |
| 7 | Chipping Sparrow | *Spizella passerina* | 206 | 241 | 180 | 210 | 212 | 239 | 200 | 296 | 114 | 152 | 2050 |
| 8 | Cassin's Kingbird | *Tyrannus vociferans* | 221 | 91 | 80 | 50 | 29 | 181 | 222 | 287 | 234 | 239 | 1634 |
| 9 | Barn Swallow | *Hirundo rustica* | 160 | 117 | 110 | 115 | 103 | 150 | 112 | 82 | 105 | 275 | 1329 |
| 10 | House Finch | *Haemorhous mexicanus* | 192 | 181 | 70 | 50 | 51 | 84 | 112 | 114 | 103 | 66 | 1023 |
| 11 | Northern Mockingbird | *Mimus polyglottos* | 99 | 38 | 50 | 80 | 70 | 42 | 57 | 101 | 82 | 121 | 740 |
| 12 | Lesser Goldfinch | *Spinus psaltria* | 85 | 26 | 32 | 50 | 46 | 69 | 106 | 70 | 135 | 100 | 719 |
| 13 | Black-throated Sparrow | *Amphispiza bilineata* | 95 | 93 | 85 | 60 | 50 | 30 | 82 | 58 | 73 | 30 | 656 |
| 14 | Cave Swallow | *Petrochelidon fulva* | 34 | 0 | 50 | 10 | 236 | 26 | 28 | 132 | 25 | 44 | 585 |
| 15 | Pyrrhuloxia | *Cardinalis sinuatus* | 44 | 67 | 35 | 55 | 56 | 35 | 48 | 58 | 80 | 89 | 567 |
| 16 | Lark Sparrow | *Chondestes grammacus* | 0 | 74 | 25 | 30 | 19 | 17 | 25 | 70 | 145 | 161 | 566 |
| 17 | Turkey Vulture | *Cathartes aura* | 37 | 54 | 50 | 58 | 54 | 74 | 46 | 64 | 87 | 38 | 562 |
| 18 | Eastern Meadowlark | *Sturnella magna* | 46 | 75 | 52 | 30 | 47 | 29 | 39 | 101 | 85 | 37 | 541 |
| 19 | Blue Grosbeak | *Passerina caerulea* | 15 | 20 | 25 | 30 | 45 | 58 | 60 | 100 | 67 | 119 | 539 |
| 20 | Bell's Vireo | *Vireo bellii* | 41 | 74 | 30 | 40 | 29 | 20 | 32 | 79 | 76 | 79 | 500 |
| 21 | Brown-headed Cowbird | *Molothrus ater* | 53 | 52 | 40 | 30 | 38 | 32 | 176 | 30 | 2 | 30 | 483 |
| 22 | Eurasian Collared Dove | *Streptopelia decaocto* | 0 | 29 | 10 | 12 | 35 | 0 | 14 | 68 | 94 | 164 | 426 |
| 23 | Cactus Wren | *Campylorhynchus brunneicapillus* | 32 | 69 | 25 | 40 | 78 | 22 | 27 | 52 | 28 | 44 | 417 |
| 24 | Killdeer | *Charadrius vociferus* | 32 | 39 | 38 | 42 | 23 | 30 | 45 | 61 | 62 | 32 | 404 |
| 25 | Mexican Duck | *Anas platyrhynchos* | 26 | 44 | 30 | 56 | 41 | 48 | 13 | 42 | 30 | 51 | 381 |
| 26 | Scaled Quail | *Callipepla squamata* | 8 | 36 | 30 | 50 | 7 | 13 | 37 | 82 | 62 | 52 | 377 |
| 27 | Red-billed Pigeon | *Patagioenas flavirostris* | 0 | 66 | 0 | 70 | 0 | 0 | 0 | 0 | 0 | 238 | 374 |
| 28 | Gold-front Woodpecker | *Melanerpes aurifrons* | 4 | 5 | 6 | 5 | 14 | 30 | 49 | 107 | 95 | 43 | 358 |
| 29 | Curve-billed Thrasher | *Toxostoma curvirostre* | 13 | 11 | 20 | 30 | 87 | 24 | 14 | 44 | 26 | 40 | 309 |
| 30 | Black Phoebe | *Sayornis nigricans* | 1 | 23 | 25 | 30 | 52 | 41 | 38 | 43 | 22 | 30 | 305 |
| 31 | Rock Pigeon | *Columba livia* | 2 | 0 | 10 | 12 | 40 | 26 | 20 | 101 | 76 | 0 | 287 |
| 32 | Burrowing Owl | *Athene cunicularia* | 31 | 35 | 20 | 21 | 18 | 27 | 28 | 33 | 13 | 23 | 249 |
| 33 | Cattle Egret | *Bubulcus ibis* | 0 | 12 | 20 | 28 | 4 | 0 | 18 | 39 | 111 | 14 | 246 |
| 34 | Cassin's Sparrow | *Peucaea cassinii* | 44 | 61 | 32 | 22 | 37 | 19 | 10 | 10 | 0 | 0 | 235 |
| 35 | N Rgh-wing Swallow | *Stelgidopteryx serripennis* | 2 | 40 | 10 | 10 | 0 | 88 | 22 | 0 | 15 | 12 | 199 |
| 36 | Northern Cardinal | *Cardinalis cardinalis* | 28 | 46 | 30 | 10 | 28 | 5 | 0 | 14 | 26 | 7 | 194 |
| 37 | Black Vulture | *Coragyps atratus* | 0 | 0 | 20 | 22 | 37 | 26 | 22 | 24 | 19 | 20 | 190 |
| 38 | Ladder-back Woodpkr | *Dryobates scalaris* | 12 | 24 | 6 | 14 | 8 | 22 | 14 | 36 | 29 | 23 | 188 |
| 39 | Greater Roadrunner | *Geococcyx californianus* | 19 | 17 | 18 | 16 | 15 | 12 | 19 | 28 | 27 | 17 | 188 |
| 40 | Chihuahuan Raven | *Corvus cryptoleucus* | 49 | 7 | 20 | 10 | 50 | 16 | 17 | 19 | 0 | 0 | 188 |
| 41 | Western Kingbird | *Tyrannus verticalis* | 7 | 2 | 10 | 0 | 103 | 0 | 49 | 0 | 10 | 0 | 181 |
| 42 | Say's Phoebe | *Sayornis saya* | 15 | 70 | 10 | 20 | 21 | 10 | 13 | 7 | 11 | 4 | 181 |
| 43 | European Starling | *Sturnus vulgaris* | 0 | 8 | 4 | 6 | 0 | 18 | 5 | 39 | 96 | 0 | 176 |
| 44 | Northern Flicker | *Colaptes auratus* | 31 | 30 | 20 | 10 | 26 | 0 | 5 | 6 | 22 | 23 | 173 |
| 45 | Verdin | *Auriparus flaviceps* | 0 | 3 | 8 | 10 | 5 | 3 | 18 | 40 | 25 | 58 | 170 |
| 46 | Bronzed Cowbird | *Molothrus aeneus* | 0 | 0 | 0 | 0 | 0 | 0 | 0 | 141 | 20 | 0 | 161 |
| 47 | Black-chinned Hummingbird | *Archilochus alexandri* | 8 | 14 | 10 | 8 | 17 | 7 | 15 | 49 | 26 | 5 | 159 |
| 48 | Ash-throat Flycatcher | *Myiarchus cinerascens* | 6 | 9 | 10 | 12 | 28 | 21 | 13 | 19 | 14 | 19 | 151 |
| 49 | Common Ground-Dove | *Columbina passerina* | 32 | 28 | 20 | 20 | 28 | 12 | 0 | 0 | 0 | 0 | 140 |
| 50 | Painted Bunting | *Passerina ciris* | 0 | 2 | 0 | 2 | 2 | 16 | 19 | 24 | 32 | 39 | 136 |
| 51 | Ruf-crowned Sparrow | *Aimophila ruficeps* | 63 | 10 | 15 | 5 | 40 | 0 | 2 | 0 | 0 | 0 | 135 |
| 52 | Lesser Nighthawk | *Chordeiles acutipennis* | 2 | 0 | 4 | 2 | 8 | 24 | 0 | 0 | 18 | 57 | 115 |
| 53 | Common Nighthawk | *Chordeiles minor* | 21 | 7 | 5 | 4 | 3 | 0 | 16 | 54 | 0 | 0 | 110 |
| 54 | Loggerhead Shrike | *Lanius ludovicianus* | 8 | 8 | 8 | 10 | 14 | 4 | 20 | 16 | 5 | 12 | 105 |
| 55 | Sandhill Crane | *Antigone canadensis* | 0 | 0 | 0 | 0 | 0 | 0 | 0 | 0 | 100 | 0 | 100 |
| 56 | Rock Dove | *Columba livia* | 16 | 0 | 0 | 0 | 0 | 0 | 0 | 0 | 80 | 0 | 96 |
| 57 | American Coot | *Fulica americana* | 6 | 0 | 8 | 10 | 12 | 11 | 11 | 25 | 1 | 9 | 93 |
| 58 | Common Raven | *Corvus corax* | 21 | 6 | 10 | 8 | 7 | 2 | 10 | 12 | 10 | 2 | 88 |
|  |  |  | 2009 | 2010 | 2011 | 2012 | 2013 | 2014 | 2015 | 2016 | 2017 | 2018 | TOTAL |
| 59 | Canyon Towhee | *Melospiza fusca* | 10 | 0 | 8 | 0 | 51 | 6 | 6 | 0 | 1 | 6 | 88 |
| 60 | Western Meadowlark | *Sturnella neglecta* | 6 | 42 | 4 | 2 | 3 | 0 | 0 | 15 | 0 | 10 | 82 |
| 61 | Ruddy Ground-Dove | *Columbina talpacoti* | 18 | 7 | 10 | 10 | 12 | 0 | 2 | 0 | 0 | 0 | 59 |
| 62 | American Kestrel | *Falco sparverius* | 11 | 2 | 5 | 6 | 9 | 10 | 6 | 9 | 0 | 0 | 58 |
| 63 | Swainson's Hawk | *Buteo swainsoni* | 5 | 3 | 5 | 6 | 7 | 6 | 5 | 5 | 9 | 6 | 57 |
| 64 | Common Yellowthroat | *Geothlypis trichas* | 0 | 0 | 0 | 0 | 0 | 8 | 5 | 12 | 8 | 14 | 47 |
| 65 | Great Blue Heron | *Ardea herodias* | 2 | 2 | 6 | 2 | 11 | 0 | 2 | 7 | 0 | 14 | 46 |
| 66 | Black-chinned Sparrow | *Spizella atrogularis* | 7 | 2 | 2 | 0 | 32 | 0 | 2 | 0 | 0 | 0 | 45 |
| 67 | Scott's Oriole | *Icterus parisorum* | 3 | 0 | 2 | 0 | 25 | 6 | 8 | 0 | 0 | 0 | 44 |
| 68 | White-tailed Kite | *Elanus leucurus* | 6 | 0 | 2 | 0 | 4 | 7 | 11 | 5 | 6 | 2 | 43 |
| 69 | Vermilion Flycatcher | *Pyrocephalus rubinus* | 0 | 1 | 0 | 4 | 0 | 0 | 13 | 0 | 11 | 14 | 43 |
| 70 | Orchard Oriole | *Icterus spurius* | 14 | 4 | 2 | 2 | 1 | 1 | 2 | 7 | 8 | 0 | 41 |
| 71 | Phainopepla | *Phainopepla nitens* | 0 | 0 | 0 | 5 | 9 | 3 | 7 | 10 | 0 | 2 | 36 |
| 72 | Green Heron | *Butorides virescens* | 4 | 4 | 2 | 0 | 0 | 10 | 0 | 6 | 3 | 6 | 35 |
| 73 | Yellow-headed Black Brd | *Xanthocephalus xanthocephalus* | 10 | 3 | 0 | 0 | 0 | 0 | 0 | 0 | 0 | 20 | 33 |
| 74 | Rock Wren | *Salpinctes obsoletus* | 7 | 0 | 5 | 0 | 13 | 3 | 5 | 0 | 0 | 0 | 33 |
| 75 | Red-shafted Flicker | *Colaptes auratus* | 2 | 2 | 4 | 0 | 0 | 15 | 10 | 0 | 0 | 0 | 33 |
| 76 | White-Faced Ibis | *Plegadis chihi* | 0 | 0 | 0 | 6 | 0 | 0 | 16 | 0 | 0 | 10 | 32 |
| 77 | Northern Bobwhite | *Colinus virginianus* | 3 | 5 | 4 | 2 | 5 | 3 | 6 | 0 | 4 | 0 | 32 |
| 78 | Whip-poor-will | *Antrostomus vociferus* | 5 | 3 | 2 | 3 | 8 | 2 | 0 | 0 | 0 | 5 | 28 |
| 79 | Grasshopper Sparrow | *Ammodramus savannarum* | 2 | 0 | 2 | 0 | 7 | 11 | 0 | 0 | 6 | 0 | 28 |
| 80 | Yellow-Billed Cuckoo | *Coccyzus americanus* | 0 | 0 | 0 | 0 | 0 | 0 | 4 | 12 | 9 | 2 | 27 |
| 81 | Red-tailed Hawk | *Buteo jamaicensis* | 1 | 4 | 2 | 3 | 2 | 4 | 9 | 0 | 2 | 0 | 27 |
| 82 | Fulvous Whistl Duck | *Dendrocygna bicolor* | 0 | 2 | 0 | 1 | 17 | 5 | 0 | 0 | 0 | 0 | 25 |
| 83 | Horned Lark | *Eremophila alpestris* | 0 | 2 | 1 | 1 | 3 | 6 | 6 | 3 | 1 | 0 | 23 |
| 84 | Indigo Bunting | *Passerina cyanea* | 3 | 5 | 2 | 2 | 7 | 0 | 1 | 0 | 1 | 0 | 21 |
| 85 | Hooded Oriole | *Icterus cucullatus* | 6 | 4 | 0 | 1 | 0 | 0 | 3 | 4 | 2 | 0 | 20 |
| 86 | Black-crowned Night Heron | *Nycticorax nycticorax* | 0 | 0 | 0 | 0 | 0 | 0 | 0 | 0 | 20 | 0 | 20 |
| 87 | Yellow-breasted Chat | *Icteria virens* | 0 | 0 | 0 | 1 | 1 | 0 | 2 | 7 | 5 | 3 | 19 |
| 88 | Blue-gray Gnatcatcher | *Poliophtila caerulea* | 0 | 0 | 0 | 0 | 0 | 0 | 0 | 2 | 7 | 9 | 18 |
| 89 | Lucifer Hummingbird | *Calothorax lucifer* | 1 | 1 | 1 | 0 | 1 | 0 | 0 | 9 | 0 | 2 | 15 |
| 90 | Common Moorhen | *Gallinula galeata* | 0 | 1 | 0 | 0 | 0 | 0 | 0 | 0 | 10 | 4 | 15 |
| 91 | Aztec Rail | *Rallus tenuirostris* | 9 | 3 | 1 | 0 | 1 | 0 | 0 | 0 | 0 | 0 | 14 |
| 92 | Varied Bunting | *Passerina versicolor* | 2 | 1 | 0 | 0 | 5 | 0 | 0 | 0 | 3 | 3 | 14 |
| 93 | Great Horned Owl | *Bubo virginianus* | 1 | 2 | 0 | 0 | 0 | 1 | 1 | 8 | 0 | 1 | 14 |
| 94 | Gila Woodpecker | *Melanerpes uropygialis* | 12 | 0 | 1 | 0 | 0 | 1 | 0 | 0 | 0 | 0 | 14 |
| 95 | Common Pauraque | *Nyctidromus albigollis* | 6 | 0 | 4 | 3 | 1 | 0 | 0 | 0 | 0 | 0 | 14 |
| 96 | Ruddy Duck | *Oxyura jamaicensis* | 3 | 4 | 1 | 0 | 0 | 3 | 2 | 0 | 0 | 0 | 13 |
| 97 | Black-headed Grosbeak | *Pheucticus melanocephalus* | 3 | 2 | 2 | 0 | 6 | 0 | 0 | 0 | 0 | 0 | 13 |
| 98 | Couch's Kingbird | *Tyrannus couchii* | 0 | 5 | 1 | 1 | 3 | 1 | 0 | 1 | 0 | 0 | 12 |
| 99 | Savannah Sparrow | *Passerculus sandwichensis* | 0 | 0 | 0 | 1 | 1 | 0 | 4 | 0 | 2 | 0 | 8 |
| 100 | Harris's Hawk | *Parabuteo unicinctus* | 0 | 0 | 0 | 0 | 0 | 0 | 6 | 0 | 0 | 0 | 6 |
| 101 | Bullock's Oriole | *Icterus bullockii* | 0 | 0 | 0 | 0 | 0 | 0 | 2 | 0 | 0 | 4 | 6 |
| 102 | Eastern Phoebe | *Sayornis phoebe* | 3 | 0 | 0 | 0 | 0 | 0 | 0 | 0 | 0 | 2 | 5 |
| 103 | White-crowned Sparrow | *Zonotrichia leucophrys* | 2 | 0 | 0 | 0 | 0 | 0 | 2 | 0 | 0 | 0 | 4 |
| 104 | Summer Tanager | *Piranga rubra* | 1 | 0 | 0 | 0 | 0 | 0 | 0 | 0 | 2 | 1 | 4 |
| 105 | Mississippi Kite | *Ictinia mississippiensis* | 0 | 0 | 0 | 0 | 0 | 0 | 0 | 2 | 2 | 0 | 4 |
| 106 | Clapper Rail | *Rallus crepitans* | 0 | 0 | 0 | 0 | 0 | 0 | 0 | 4 | 0 | 0 | 4 |
| 107 | Blue-winged Teal | *Spatula discors* | 0 | 4 | 0 | 0 | 0 | 0 | 0 | 0 | 0 | 0 | 4 |
| 108 | Black-tailed Gnatcatcher | *Poliophtila melanura* | 4 | 0 | 0 | 0 | 0 | 0 | 0 | 0 | 0 | 0 | 4 |
| 109 | Sora | *Porzana carolina* | 2 | 0 | 0 | 0 | 0 | 0 | 0 | 1 | 0 | 0 | 3 |
| 110 | Purple Gallinule | *Porphyrio martinicus* | 0 | 0 | 0 | 0 | 0 | 3 | 0 | 0 | 0 | 0 | 3 |
|  |  |  | 2009 | 2010 | 2011 | 2012 | 2013 | 2014 | 2015 | 2016 | 2017 | 2018 | TOTAL |
| 111 | Belted Kingfisher | *Megaceryle alcyon* | 1 | 1 | 0 | 0 | 0 | 0 | 0 | 0 | 1 | 0 | 3 |
| 112 | Altamira Oriole | *Icterus gularis* | 3 | 0 | 0 | 0 | 0 | 0 | 0 | 0 | 0 | 0 | 3 |
| 113 | Yellow Warbler | *Setophaga petechia* | 0 | 0 | 0 | 0 | 2 | 0 | 0 | 0 | 0 | 0 | 2 |
| 114 | White-tipped Dove | *Leptotila verreauxi* | 2 | 0 | 0 | 0 | 0 | 0 | 0 | 0 | 0 | 0 | 2 |
| 115 | Pied-billed Grebe | *Podilymbus podiceps* | 2 | 0 | 0 | 0 | 0 | 0 | 0 | 0 | 0 | 0 | 2 |
| 116 | Great Egret | *Ardea alba* | 0 | 0 | 0 | 0 | 0 | 0 | 0 | 1 | 1 | 0 | 2 |
| 117 | Elf Owl | *Micrathene whitneyi* | 0 | 2 | 0 | 0 | 0 | 0 | 0 | 0 | 0 | 0 | 2 |
| 118 | Cinnamon Teal | *Spatula cyanoptera* | 0 | 0 | 0 | 0 | 2 | 0 | 0 | 0 | 0 | 0 | 2 |
| 119 | Black-necked Stilt | *Himantopus mexicanus* | 0 | 0 | 0 | 0 | 0 | 2 | 0 | 0 | 0 | 0 | 2 |
| 120 | Black-bellied Whistling-Duck | *Dendrocygna autumnalis* | 0 | 0 | 0 | 0 | 0 | 0 | 0 | 0 | 2 | 0 | 2 |
| 121 | Barn Owl | *Tyto alba* | 0 | 0 | 0 | 0 | 0 | 0 | 0 | 0 | 2 | 0 | 2 |
| 122 | American Dipper | *Cinclus mexicanus* | 2 | 0 | 0 | 0 | 0 | 0 | 0 | 0 | 0 | 0 | 2 |
| 123 | Yellow-rumped Warbler | *Setophaga coronata* | 0 | 1 | 0 | 0 | 0 | 0 | 0 | 0 | 0 | 0 | 1 |
| 124 | Yellow-crowned Night Heron | *Nyctanassa violacea* | 0 | 0 | 0 | 0 | 0 | 0 | 0 | 0 | 0 | 1 | 1 |
| 125 | Yellow House Finch | *Haemorhous mexicanus* | 1 | 0 | 0 | 0 | 0 | 0 | 0 | 0 | 0 | 0 | 1 |
| 126 | White-throated Swift | *Aeronautes saxatalis* | 0 | 0 | 0 | 0 | 0 | 0 | 0 | 0 | 1 | 0 | 1 |
| 127 | White-tailed Hawk | *Geranoaetus albicaudatus* | 0 | 0 | 0 | 0 | 0 | 0 | 1 | 0 | 0 | 0 | 1 |
| 128 | Prairie Falcon | *Falco mexicanus* | 1 | 0 | 0 | 0 | 0 | 0 | 0 | 0 | 0 | 0 | 1 |
| 129 | Northern Pygmy-Owl | *Glaucidium gnoma* | 0 | 0 | 0 | 0 | 0 | 0 | 1 | 0 | 0 | 0 | 1 |
| 130 | Northern Harrier | *Circus hudsonius* | 0 | 0 | 0 | 0 | 0 | 0 | 1 | 0 | 0 | 0 | 1 |
| 131 | Least Flycatcher | *Empidonax minimus* | 0 | 0 | 0 | 0 | 0 | 0 | 0 | 0 | 1 | 0 | 1 |
| 132 | Lazuli Bunting | *Passerina amoena* | 0 | 0 | 0 | 0 | 1 | 0 | 0 | 0 | 0 | 0 | 1 |
| 133 | Lark Bunting | *Calamospiza melanocorys* | 1 | 0 | 0 | 0 | 0 | 0 | 0 | 0 | 0 | 0 | 1 |
| 134 | Flame-colored Tanager | *Piranga bidentate* | 1 | 0 | 0 | 0 | 0 | 0 | 0 | 0 | 0 | 0 | 1 |
| 135 | Crissal Thrasher | *Toxostoma crissale* | 1 | 0 | 0 | 0 | 0 | 0 | 0 | 0 | 0 | 0 | 1 |
| 136 | Common Black Hawk | *Buteogallus anthracinus* | 0 | 0 | 0 | 0 | 0 | 0 | 0 | 0 | 1 | 0 | 1 |
| 137 | Brown-crested Flycatcher | *Myiarchus tyrannulus* | 0 | 0 | 0 | 0 | 0 | 1 | 0 | 0 | 0 | 0 | 1 |
| 138 | American Pipit | *Anthus rubescens* | 0 | 0 | 0 | 0 | 0 | 1 | 0 | 0 | 0 | 0 | 1 |
| 139 | American Bittern | *Botaurus lentiginosus* | 0 | 0 | 0 | 0 | 0 | 1 | 0 | 0 | 0 | 0 | 1 |

From the data shown in Table 1, it was possible to graphically represent species’ behavior in both routes. Hence, Figure 2 shows a continuous increase in the total observations, except for the years 2010 and 2016; a period in which the results were more representative in terms of growth.

**Fig 2.**
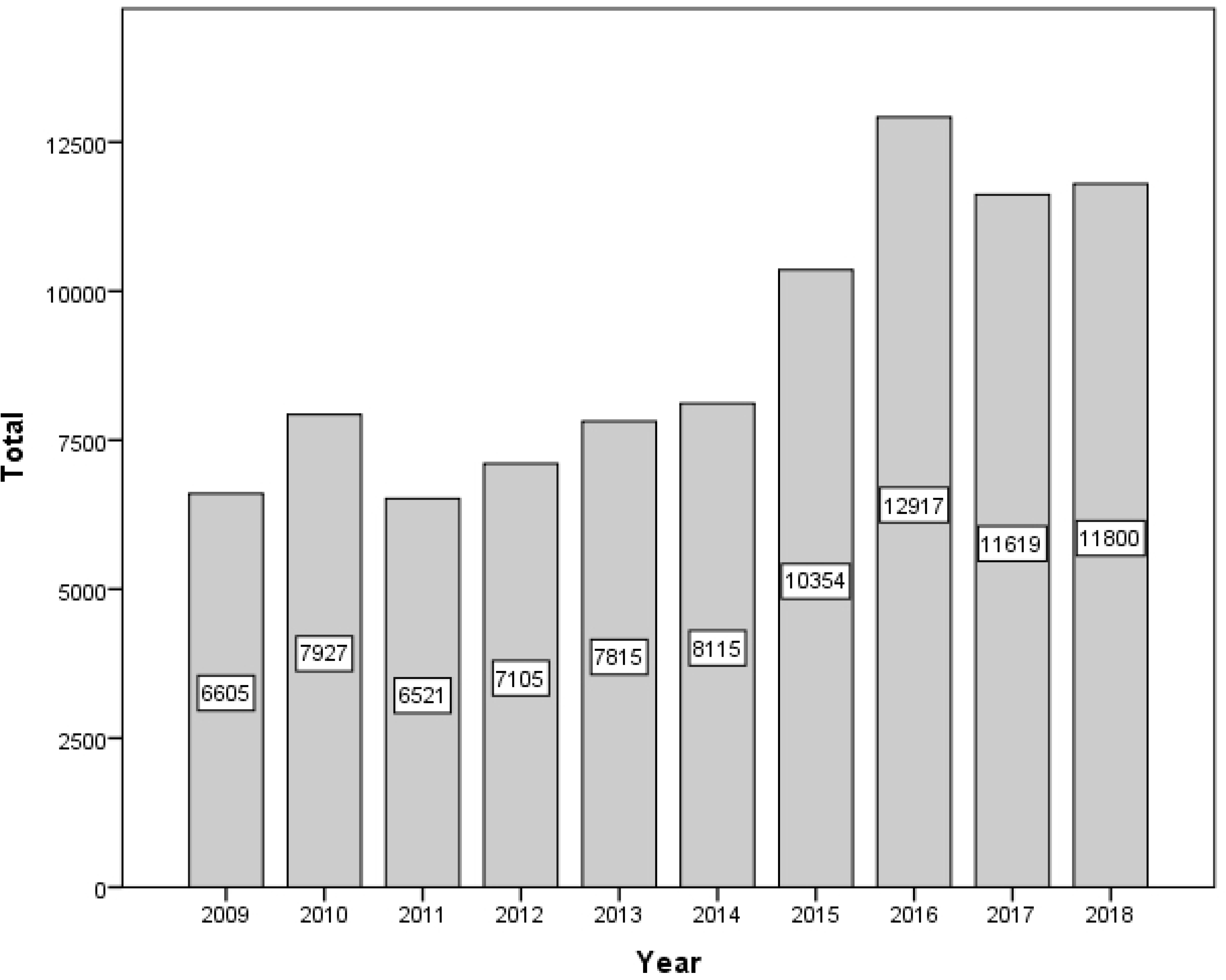
Longitudinal behavior of global data in both routes

Figure 3 shows the corresponding dispersion. The data exhibit a rising behavior as the years pass by.

**Fig 3.**
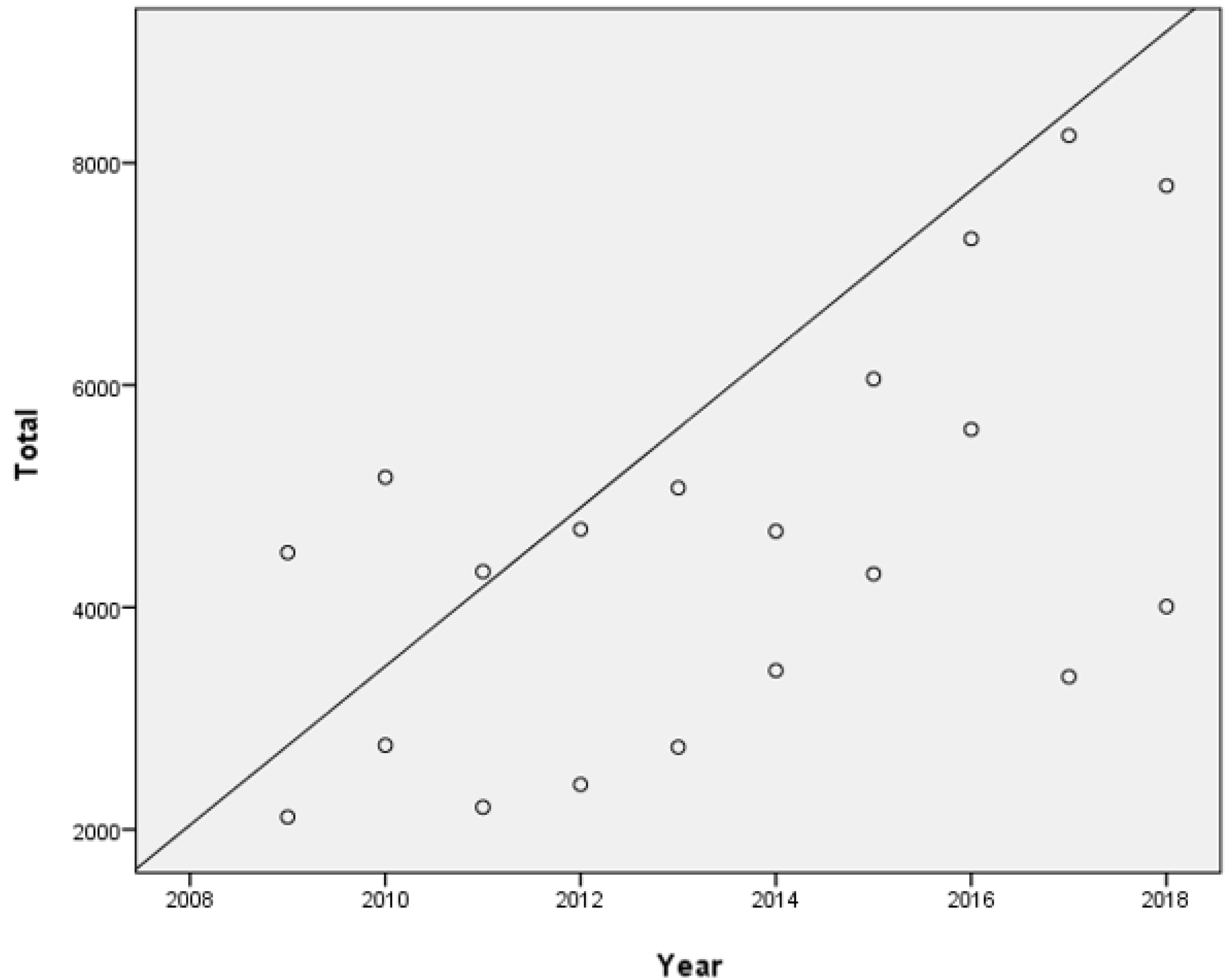
Dispersion of data.

The Pearson correlation, taking into account both routes and between the variables ‘year’ and ‘total sightings’, was of r=.569. This indicates a low correlation level. However, the significance was of .009, which corresponds to a high significance (see Table 2).

**Table 2.** Correlation of the variables ‘year’ and ‘total results’.

|  |  | Year | Total sightings |
| --- | --- | --- | --- |
| Year | Pearson Correlation | 1 | .569** |
|  | Sig. (2-tailed) |  | .009 |
|  | N | 20 | 20 |
| Total sightings | Pearson Correlation | .569** | 1 |
|  | Sig. (2-tailed) | .009 |  |
|  | N | 20 | 20 |

\*\* Correlation is significant at the 0.01 level (2-tailed).

At the moment of analyzing the data by route, we observed a similar behavior regarding the increase of the total number of sightings per year (see Figure 4).

**Fig 4.**
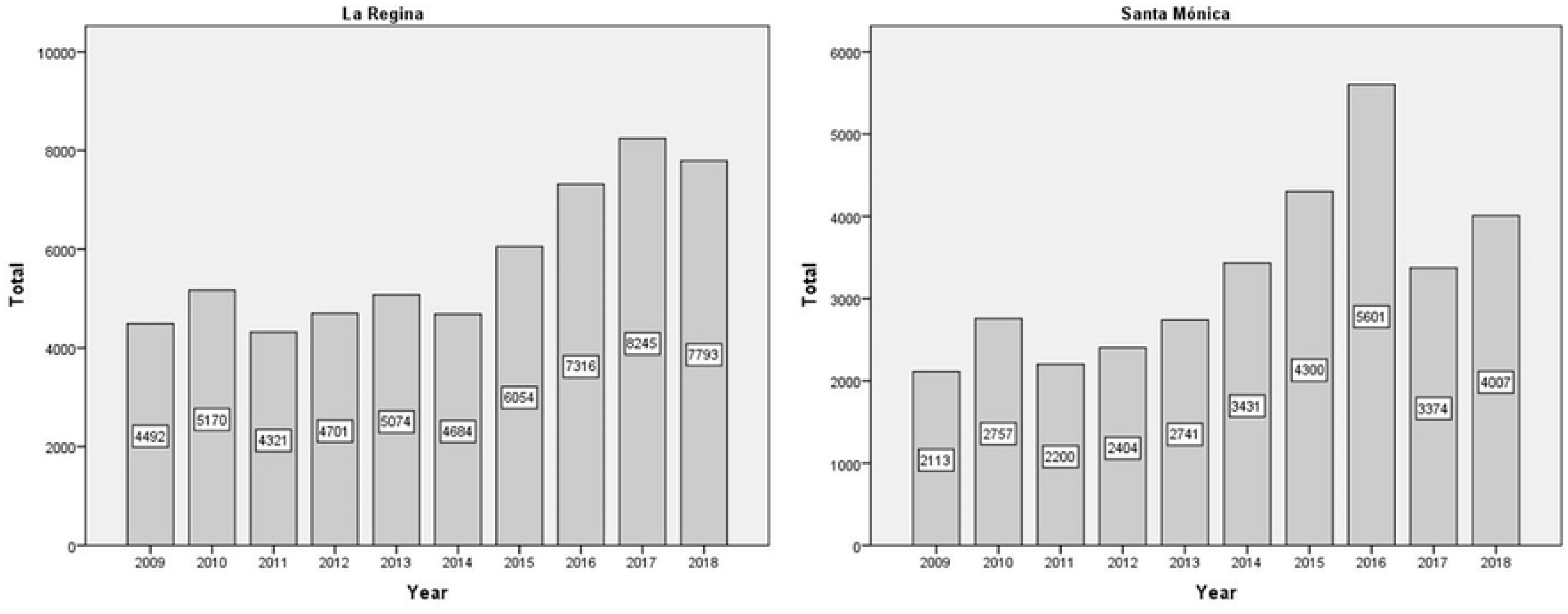
Comparison of longitudinal behaviors between both routes

When correlating the variables of total sightings against year, La Regina route shows a Sig. of .001, with a correlation coefficient r=.869 (very high) and for Santa Mónica, Sig. is .013 and its correlation coefficient is of r=.748 (high). These data are shown in Table 3.

**Table 3.**
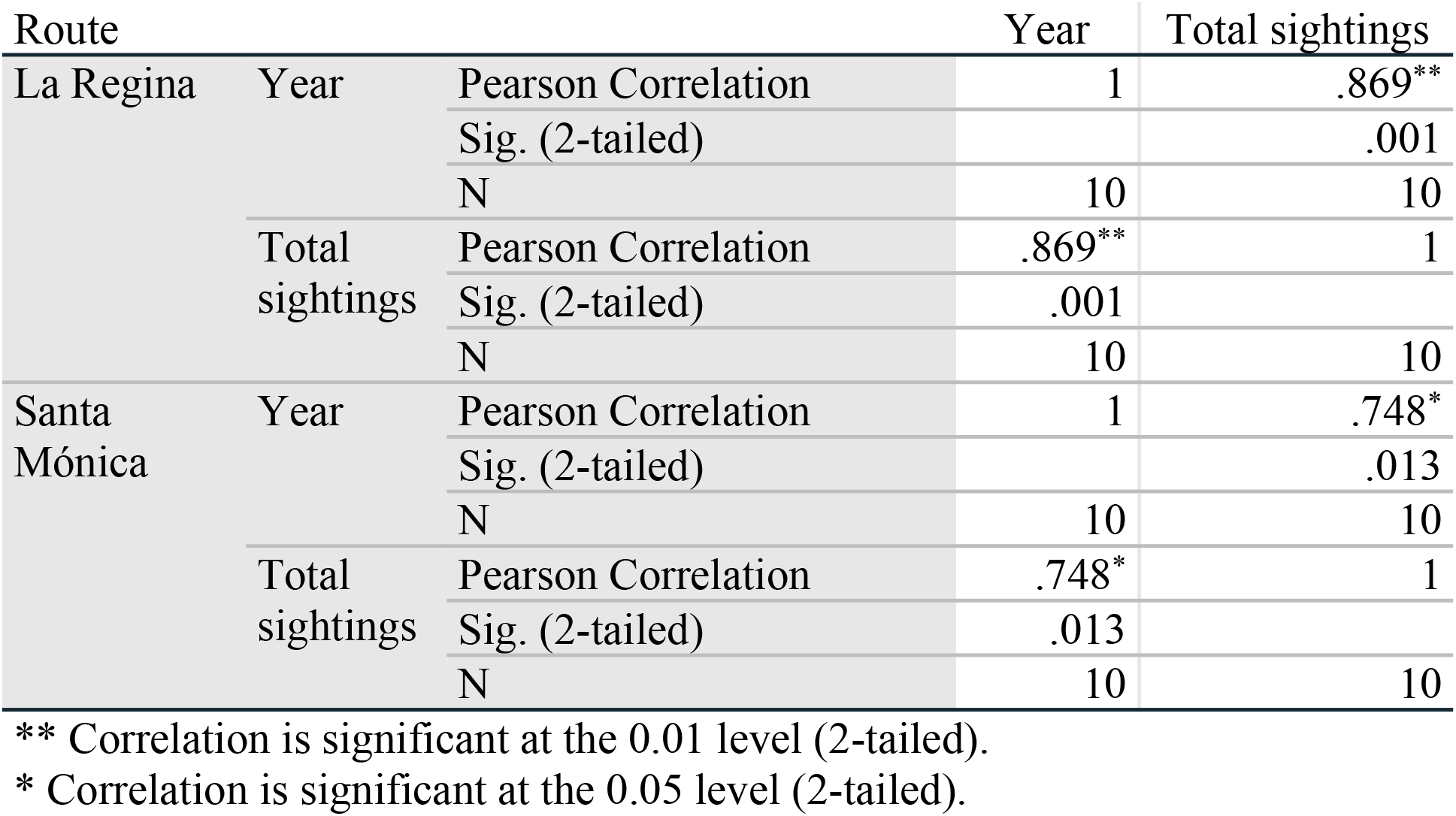
Correlations between both observation routes.

### 3.1. Increase and decrease of species during the evaluated period

The significance and the correlation analysis conducted among each year and with all the species, presented significant relations with 30 species. Table 4 shows the behavior of the species divided by route. In the cases where the correlation value is negative, it means that species decreased in their numbers and vice versa.

**Table 4.** Behavior of the species divided by routes.

| La Regina |  |  |  |  | Santa Mónica |  |  |  |
| --- | --- | --- | --- | --- | --- | --- | --- | --- |
| Species | Pearson Correlation | Sig. (2-tailed) | Significance | Correlation | Pearson Correlation | Sig. (2-tailed) | Significance | Correlation |
| *White winged dove* | .887** | 0.001 | Highly Significant | Positive, very high | .861** | 0.001 | Highly Significant | Positive, very high |
| *Mourning dove* | .845** | 0.002 | Highly Significant | Positive, very high | .940** | 0 | Highly Significant | Positive, very high |
| *Inca dove* | .793** | 0.006 | Highly Significant | Positive, high | .845** | 0.002 | Highly Significant | Positive, very high |
| *Common ground dove* | -.915** | 0 | Highly Significant | Negative, very high | -.887** | 0.001 | Highly Significant | Negative, very high |
| *Ruddy ground dove* | -.847** | 0.002 | Highly Significant | Negative, very high | -.710* | 0.021 | Significant | Negative, high |
| *Yellow billed cuckoo* | .642* | 0.045 | Significant | Positive, high | .706* | 0.022 | Significant | Positive, high |
| *Common pauraque* | -.710* | 0.021 | Significant | Negative, high | -0.058 | 0.873 |  |  |
| *Gold front woodpecker* | .786** | 0.007 | Highly Significant | Positive, high | 0.581 | 0.078 |  |  |
| *Red shafted flicker* | -0.025 | 0.946 |  |  | -.639* | 0.047 | Significant | Negative, high |
| *Vermilion flycatcher* | .688* | 0.028 | Significant | Positive, high | .640* | 0.046 | Significant | Positive, high |
| *Chihuahuan raven* | -0.526 | 0.118 |  |  | -.663* | 0.037 | Significant | Negative, high |
| *Barn swallow* | 0.284 | 0.426 |  |  | .671* | 0.034 | Significant | Positive, high |
| *Curve billed thrasher* | 0.29 | 0.416 |  |  | .814** | 0.004 | Highly Significant | Positive, very high |
| *Yellow breasted chat* | .749* | 0.013 | Significant | Positive, high | 0.607 | 0.063 |  |  |
| *Cassins sparrow* | -.909** | 0 | Highly Significant | Negative, very high | -0.455 | 0.187 |  |  |
| *Ruf crowned sparrow* | -.654* | 0.04 | Significant | Negative, high | -.863** | 0.001 | Highly Significant | Negative, very high |
| *Blk chinned sparrow* | -0.209 | 0.562 |  |  | -.817** | 0.004 | Highly Significant | Negative, very high |
| *Lark sparrow* | .708* | 0.022 | Significant | Positive, high | .747* | 0.013 | Significant | Positive, high |
| *Blk throated sparrow* | -0.622 | 0.055 |  |  | .774** | 0.009 | Highly Significant | Positive, high |
| *Northern cardinal* | -0.589 | 0.073 |  |  | -.765** | 0.01 | Highly Significant | Negative, high |
| *Blue grosbeak* | .926** | 0 | Highly Significant | Positive, very high | .740* | 0.014 | Significant | Positive, high |
| *Indigo bunting* | -0.59 | 0.072 |  |  | -.712* | 0.021 | Significant | Negative, high |
| *Painted bunting* | .942** | 0 | Highly Significant | Positive, very high | 0.399 | 0.253 |  |  |
| *Red winged blackbird* | .906** | 0 | Highly Significant | Positive, very high | .910** | 0 | Highly Significant | Positive, very high |
| *Brn headed cowbird* | -0.068 | 0.851 |  |  | .763* | 0.01 | Significant | Positive, high |
| *Lesser goldfinch* | .672* | 0.033 | Significant | Positive, high | 0.294 | 0.41 |  |  |
| *House sparrow* | 0.607 | 0.062 |  |  | .685* | 0.029 | Significant | Positive, high |
| *\*Virginairail* | -.699* | 0.025 | Significant | Negative, high | .a | . |  |  |
| *\*Eurasiancollaredove* | .768** | 0.01 | Highly Significant | Positive, high | .724* | 0.018 | Significant | Positive, high |
| *\*Verdin* | .831** | 0.003 | Highly Significant | Positive, very high | .714* | 0.02 | Significant | Positive, high |
| *\*Bluegraygnatcatcher* | .765* | 0.01 | Significant | Positive, high | .650* | 0.042 | Significant | Positive, high |

\*\* Correlation is significant at the 0.01 level (2-tailed).
\* Correlation is significant at the 0.05 level (2-tailed).

Although there are species that underwent relevant changes in only one of the routes, in the majority of the cases such changes happened in both routes. The species whose common names in English show a ‘*’ at the beginning indicate that they were identified in the monitoring, but they were not included in the initial observation lists. Such treatment is maintained throughout this paper.

Data presented in Table 4, allow identifying the main species to analyze for each route. In the case of La Regina, six species showed a decrease in their numbers per year and 15 species showed an increase. In Santa Mónica, eight species decreased and 16 increased (see Table 5).

**Table 5.** Register of increase or decrease of species by route.

| **La Regina route: Species that decreased in numbers** |  |  |  |  |
| --- | --- | --- | --- | --- |
| **Species** | **Pearson Correlation** | **Correlation** | **Sig. (2-tailed)** | **Significance** |
| *Common ground dove* | -.915** | Negative, very high | 0 | Highly Significant |
| *Cassins sparrow* | -.909** | Negative, very high | 0 | Highly Significant |
| *Ruddy ground dove* | -.847** | Negative, very high | 0.002 | Highly Significant |
| *Common pauraque* | -.710* | Negative, high | 0.021 | Significant |
| *\*Virginiarail* | -.699* | Negative, high | 0.025 | Significant |
| *Ruf crowned sparrow* | -.654* | Negative, high | 0.04 | Significant |
| **La Regina route: Species that increased in numbers** |  |  |  |  |
| **Species** | **Pearson Correlation** | **Correlation** | **Sig. (2-tailed)** | **Significance** |
| *Painted bunting* | .942** | Positive, very high | 0 | Highly Significant |
| *Blue grosbeak* | .926** | Positive, very high | 0 | Highly Significant |
| *Red winged blackbird* | .906** | Positive, very high | 0 | Highly Significant |
| *White winged dove* | .887** | Positive, very high | 0.001 | Highly Significant |
| *Mourning dove* | .845** | Positive, very high | 0.002 | Highly Significant |
| *\*Verdin* | .831** | Positive, very high | 0.003 | Highly Significant |
| *Inca dove* | .793** | Positive, high | 0.006 | Highly Significant |
| *Gold front woodpecker* | .786** | Positive, high | 0.007 | Highly Significant |
| *\*Eurasian collared dove* | .768** | Positive, high | 0.01 | Highly Significant |
| *\*Blue gray gnatcatcher* | .765* | Positive, high | 0.01 | Significant |
| *Yellow breasted chat* | .749* | Positive, high | 0.013 | Significant |
| *Lark sparrow* | .708* | Positive, high | 0.022 | Significant |
| *Vermilion flycatcher* | .688* | Positive, high | 0.028 | Significant |
| *Lesser goldfinch* | .672* | Positive, high | 0.033 | Significant |
| *Yellow billed cuckoo* | .642* | Positive, high | 0.045 | Significant |
| **Santa Mónica route: Species that decreased in numbers** |  |  |  |  |
| **Species** | **Pearson Correlation** | **Correlation** | **Sig. (2-tailed)** | **Significance** |
| *Common ground dove* | -.887** | Negative, very high | 0.001 | Highly Significant |
| *Ruf crowned sparrow* | -.863** | Negative, very high | 0.001 | Highly Significant |
| *Blk chinned sparrow* | -.817** | Negative, very high | 0.004 | Highly Significant |
| *Northern cardinal* | -.765** | Negative, high | 0.01 | Highly Significant |
| *Indigo bunting* | -.712* | Negative, high | 0.021 | Significant |

| *Ruddy ground dove* | -.710* | Negative, high | 0.021 | Significant |
| --- | --- | --- | --- | --- |
| *Chihuahuan raven* | -.663* | Negative, high | 0.037 | Significant |
| *Red shafted flicker* | -.639* | Negative, high | 0.047 | Significant |
| **Santa Mónica route: Species that increased in numbers** |  |  |  |  |
| **Species** | **Pearson Correlation** | **Correlation** | **Sig. (2-tailed)** | **Significance** |
| *Mourning dove* | .940** | Positive, very high | 0 | Highly Significant |
| *Red winged blackbird* | .910** | Positive, very high | 0 | Highly Significant |
| *White winged dove* | .861** | Positive, very high | 0.001 | Highly Significant |
| *Inca dove* | .845** | Positive, very high | 0.002 | Highly Significant |
| *Curve billed thrasher* | .814** | Positive, very high | 0.004 | Highly Significant |
| *Blk throated sparrow* | .774** | Positive, high | 0.009 | Highly Significant |
| *Brn headed cowbird* | .763* | Positive, high | 0.01 | Significant |
| *Lark sparrow* | .747* | Positive, high | 0.013 | Significant |
| *Blue grosbeak* | .740* | Positive, high | 0.014 | Significant |
| *\*Eurasian collared dove* | .724* | Positive, high | 0.018 | Significant |
| *\*Verdin* | .714* | Positive, high | 0.02 | Significant |
| *Yellow billed cuckoo* | .706* | Positive, high | 0.022 | Significant |
| *House sparrow* | .685* | Positive, high | 0.029 | Significant |
| *Barn swallow* | .671* | Positive, high | 0.034 | Significant |
| *\*Blue gray gnat catcher* | .650* | Positive, high | 0.042 | Significant |
| *Vermilion flycatcher* | .640* | Positive, high | 0.046 | Significant |

### 3.2. Species that decreased in numbers

In this section, we present the analysis of the changes in the sightings of species, starting with the ones that shown a dramatic decrease in their numbers, according to the Pearson correlation coefficient and significance.

**Species:** Common ground dove (*Columbina passerina*)

**Link:** https://www.naturalista.mx/taxa/3545-Columbina-passerina

**Condition:** This species might have moved some years from the zone of Parque Nacional Big Bend. From 2015, it was not observed anymore (see Figure 5).

**Fig 5.**
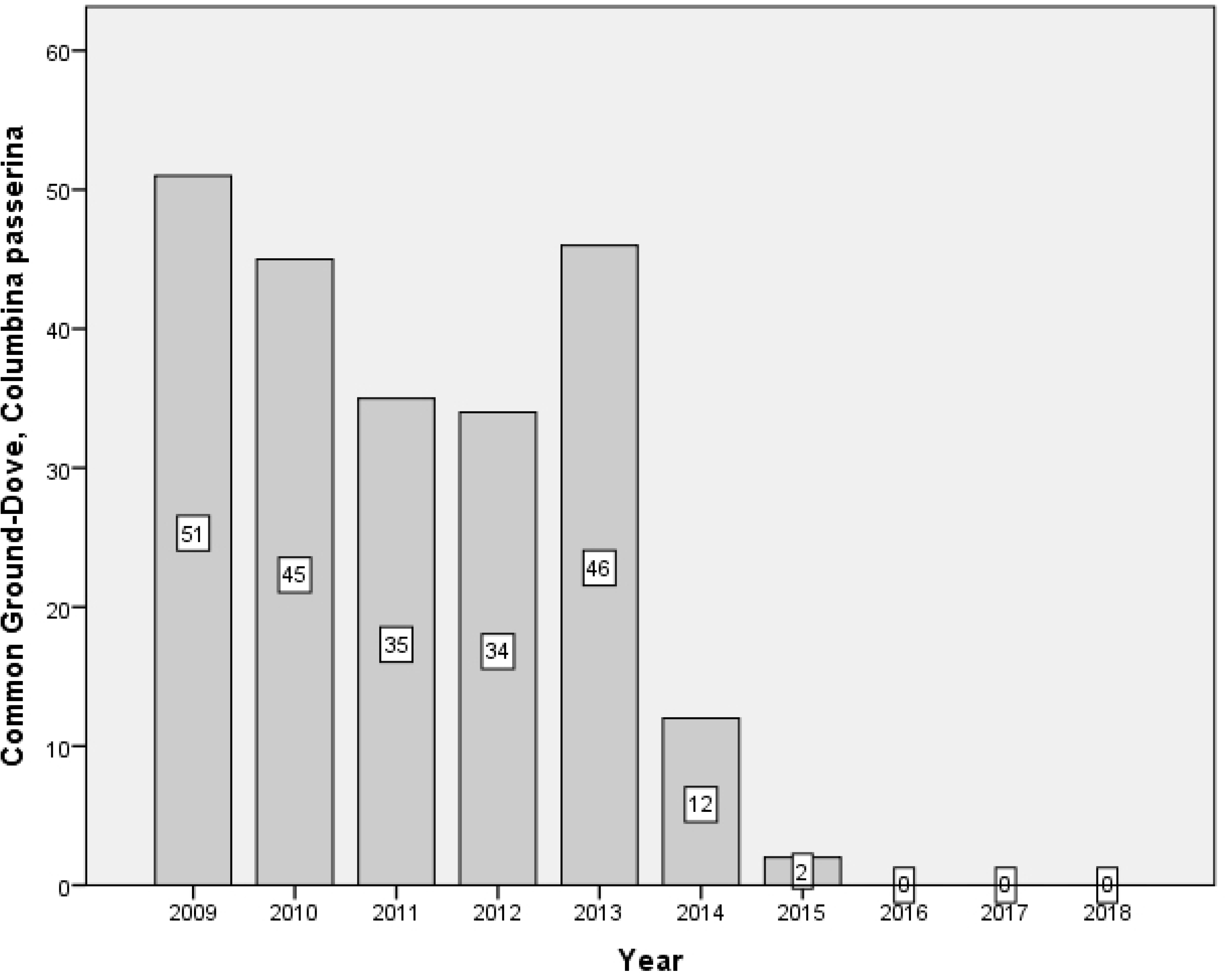
Decrease of the Common ground dove (*Columbina passerina*) species

**Species:** Ruddy ground dove (*Columbina talpacoti)*

**Link:** https://ebird.org/species/rugdov

**Condition:** This species of pigeon is not common in this zone, and it only showed up from 2009 to 2013. Normally, this bird has a permanent presence in coast areas (see Figure 6).

**Fig 6.**
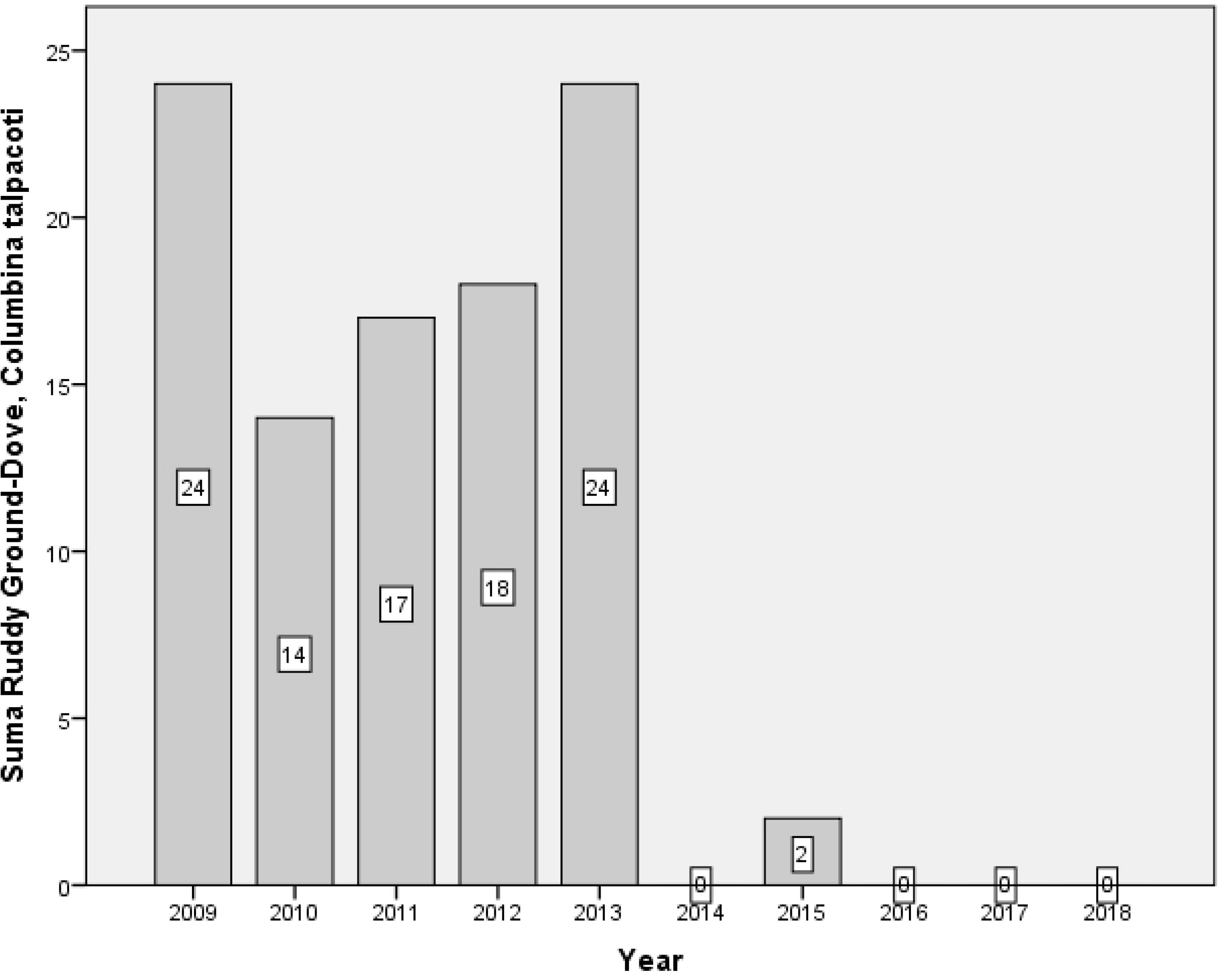
Decrease of the Ruddy ground dove (*Columbina talpacoti)* species

**Fig 7.**
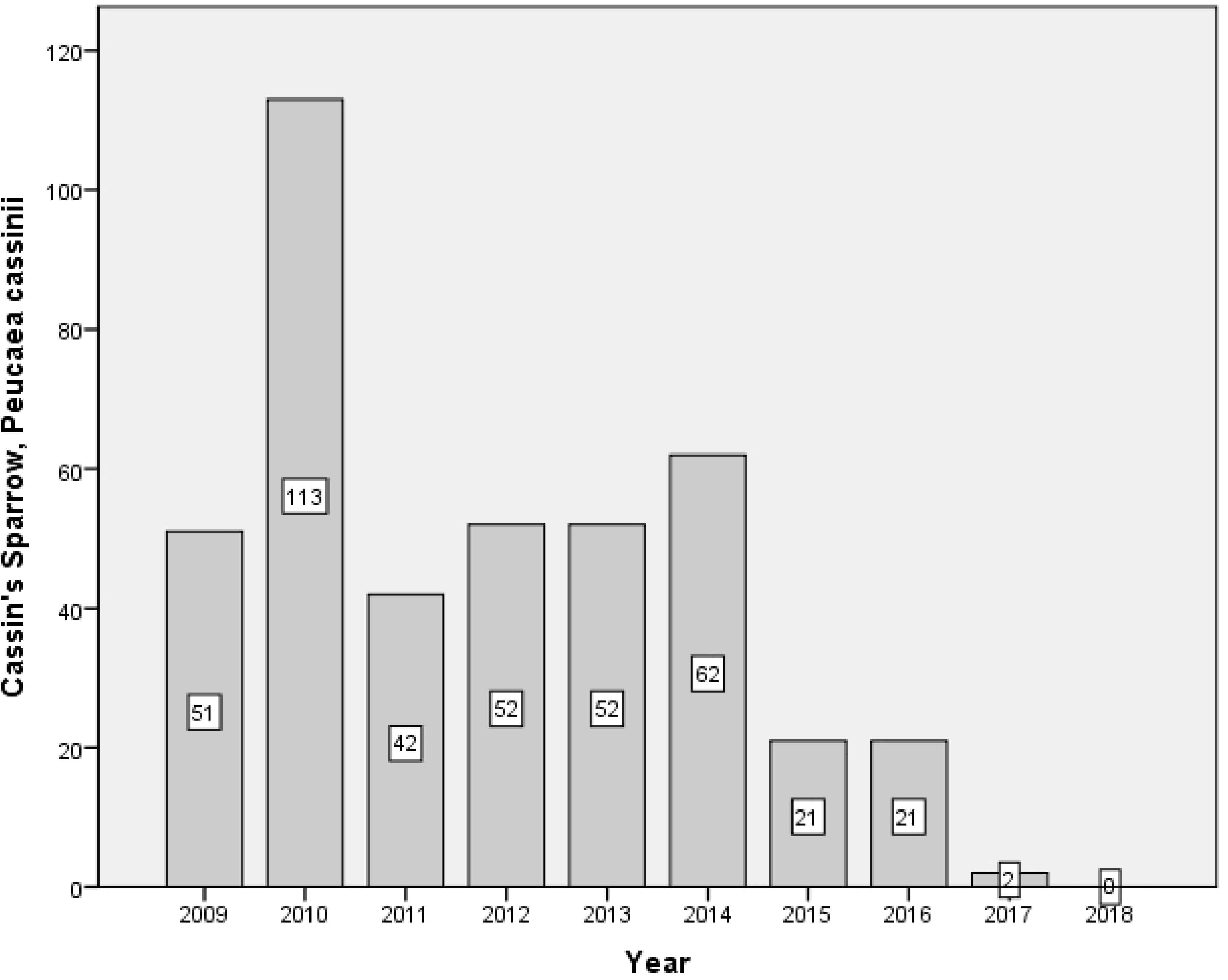
Decrease of the Cassin’s sparrow (*Peucaea cassinii)* species

**Fig 8.**
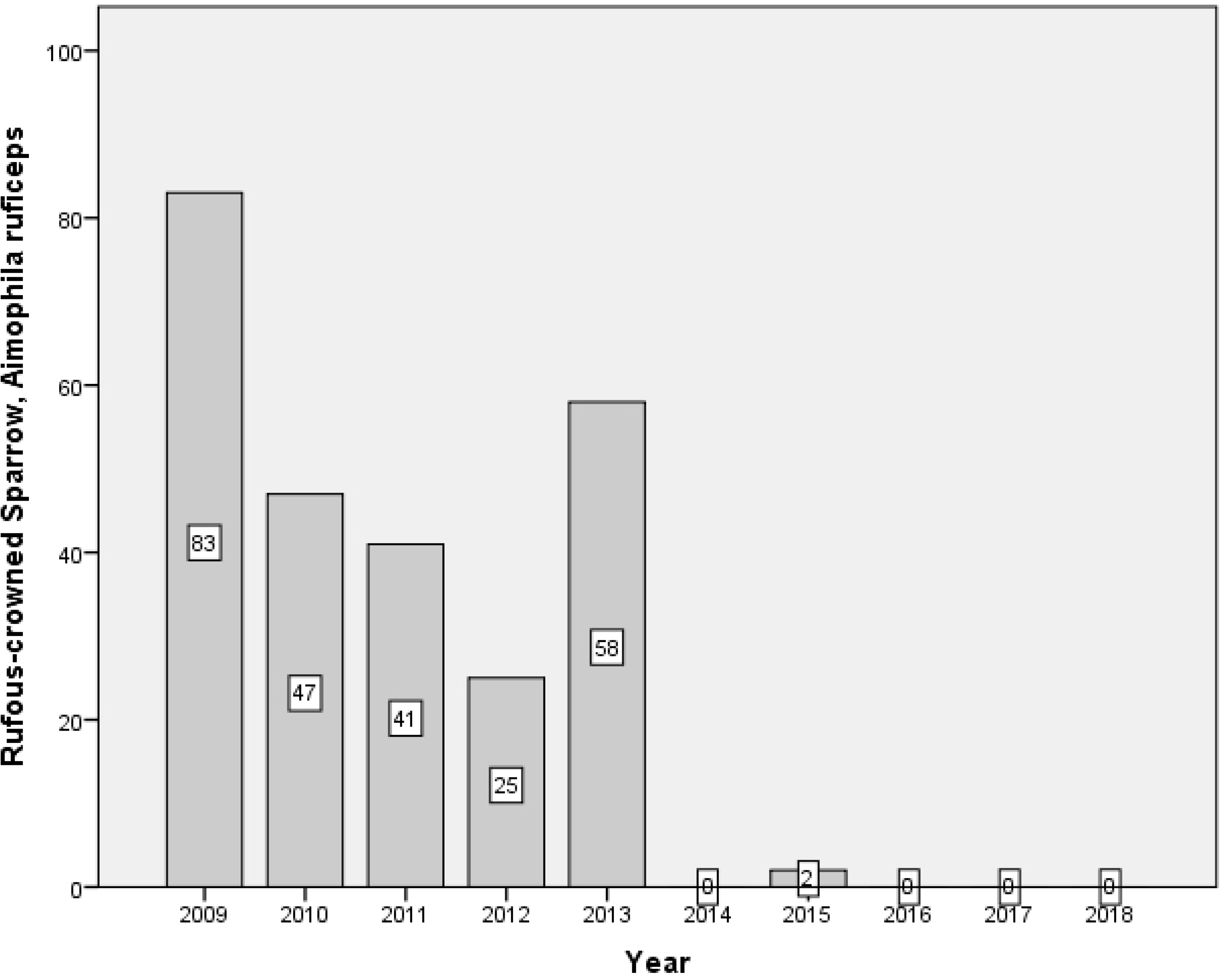
Decrease of the Rufous crowned sparrow (*Aimophila ruficeps*) species

**Fig 9.**
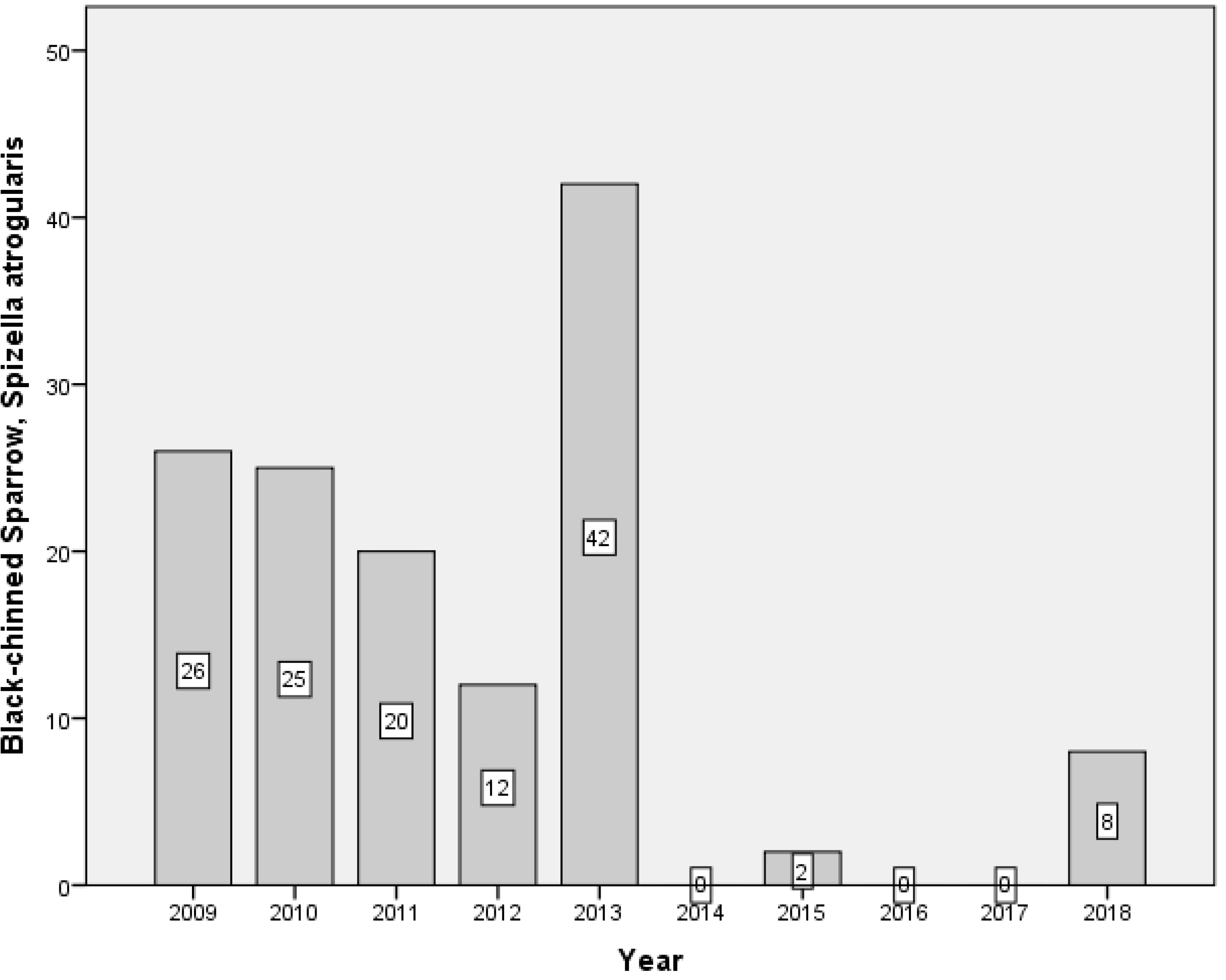
Decrease of the Black-chinned sparrow (*Spizella atrogularis*) species

**Fig 10.**
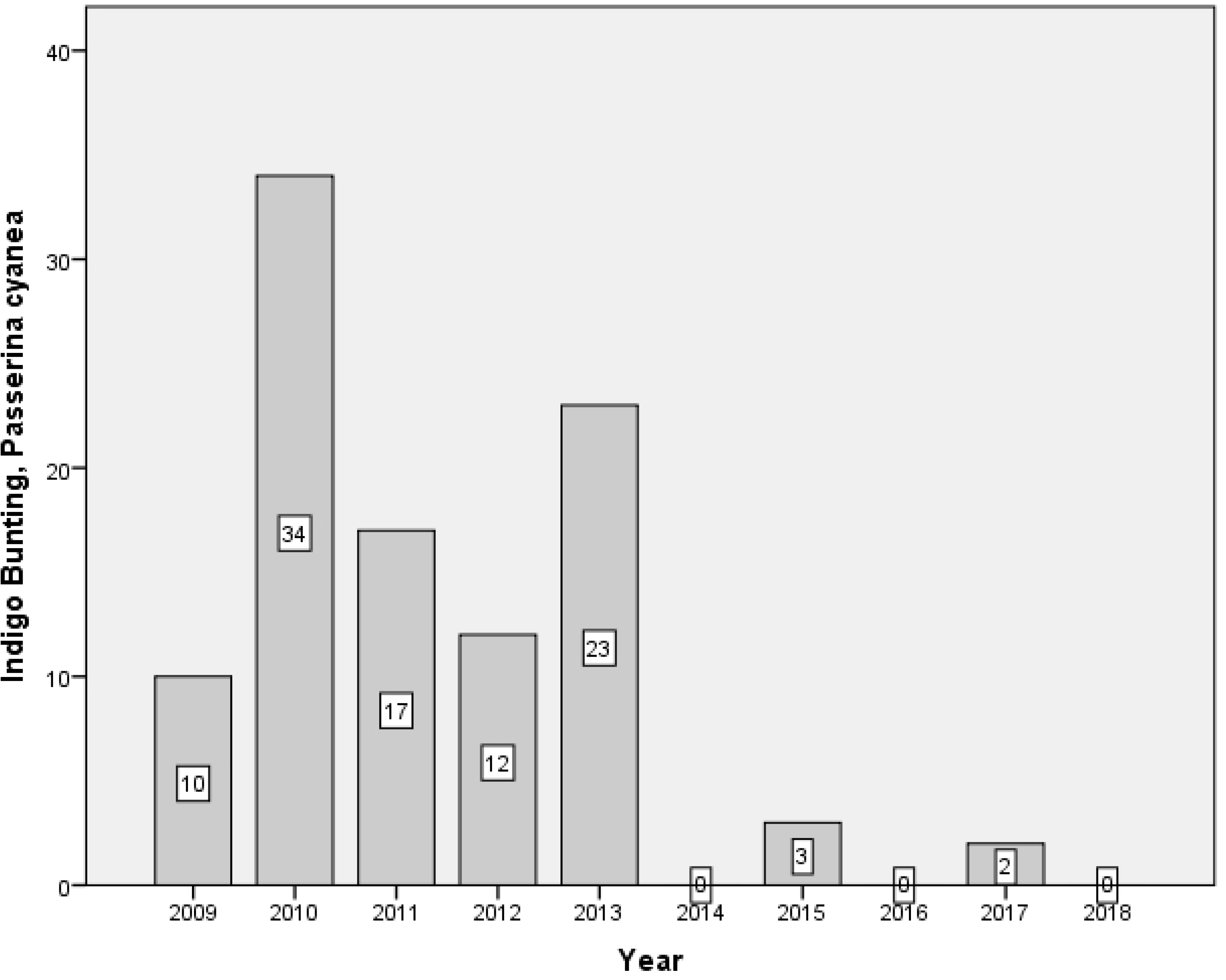
Decrease of the Indigo bunting (*Passerina cyanea)* species

**Fig 11.**
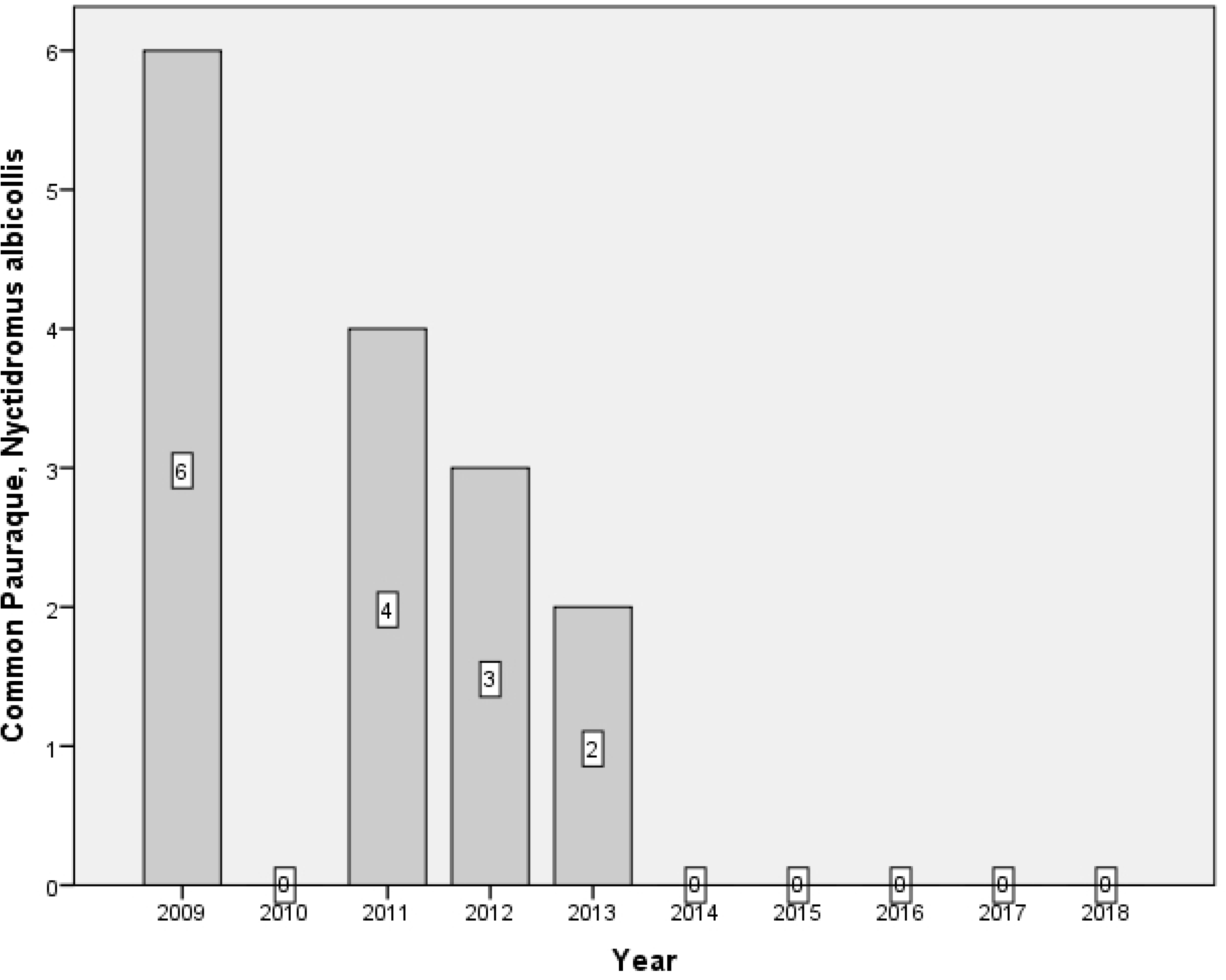
Decrease of the Common pauraque (*Nyctidromus albicollis*) species

**Fig 12.**
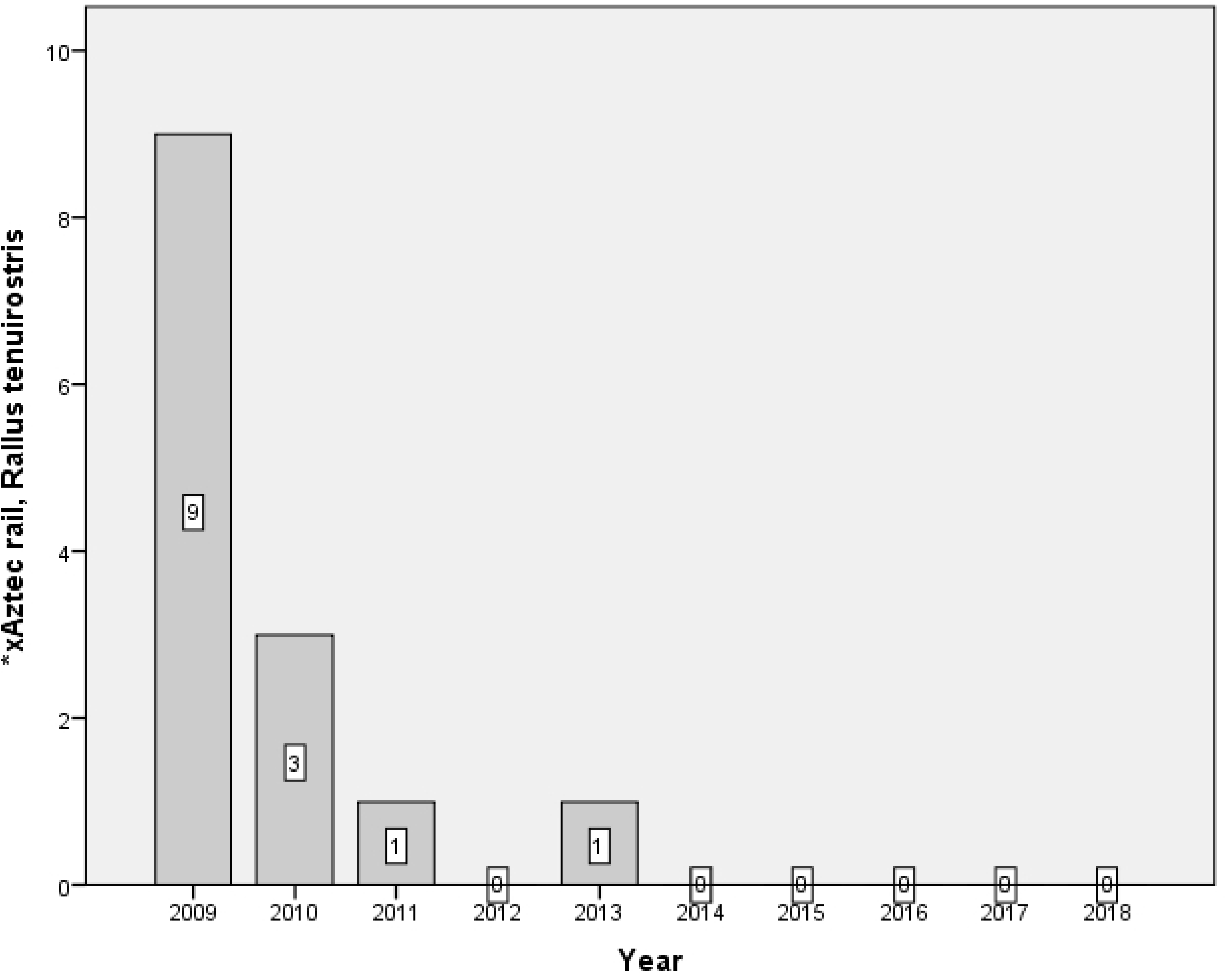
Decrease of the *Aztec rail (*Rallus tenuirostris)* species

**Fig 13.**
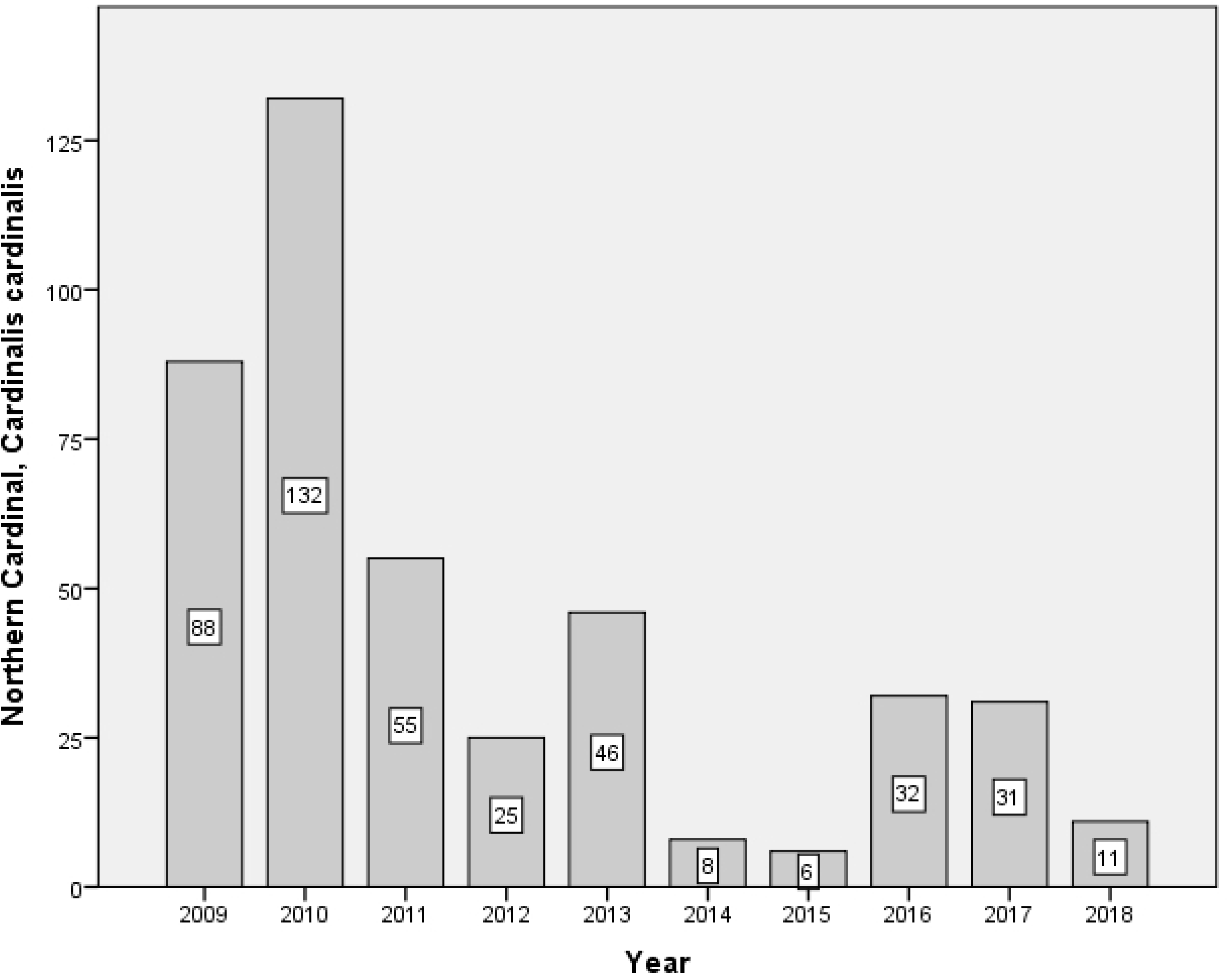
Conditions of the Northern cardinal (*Cardinalis cardinalis)* species

**Fig 14.**
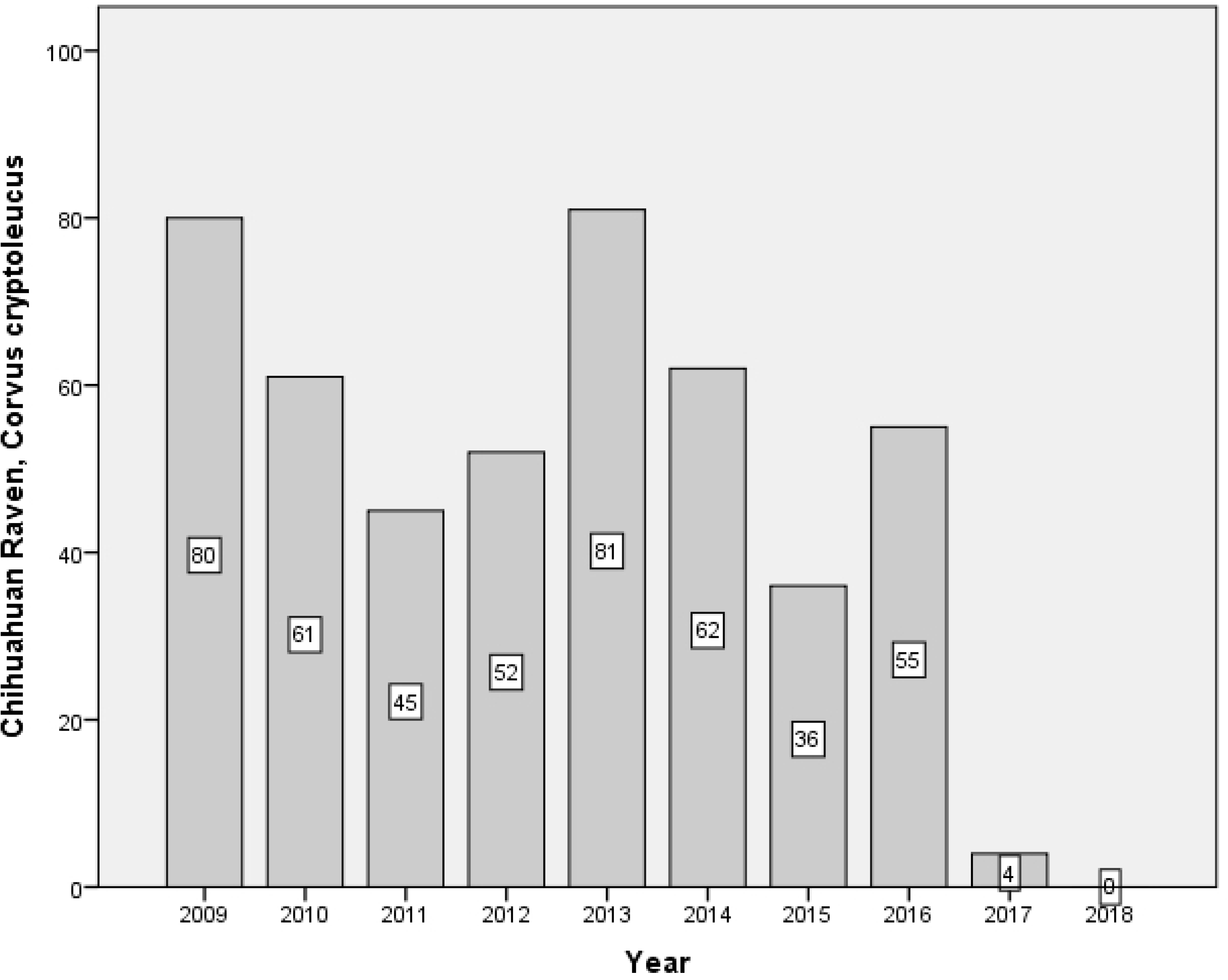
Conditions of the Chihuahua raven (*Corvus cryptoleucus*) species

**Fig 15.**
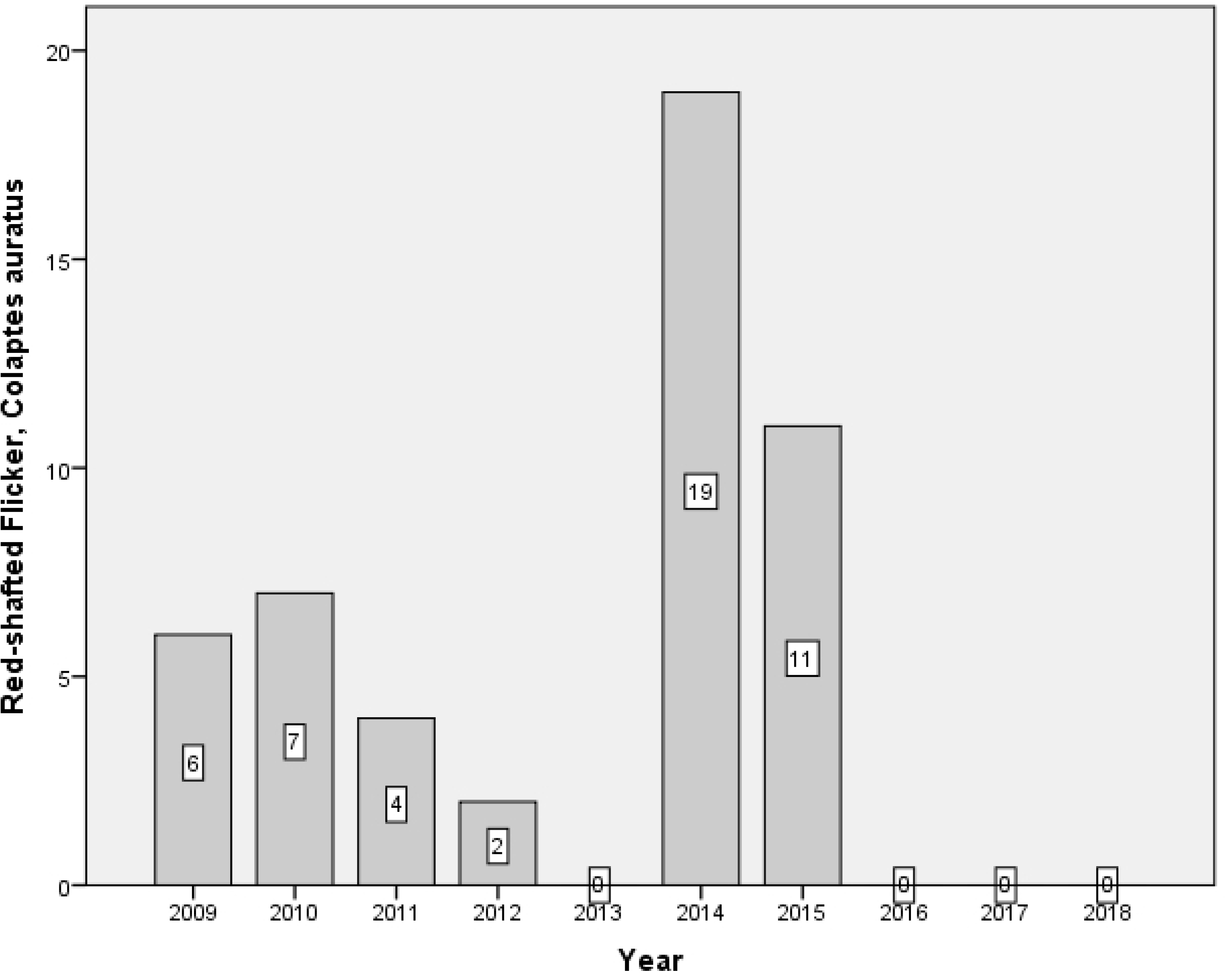
Conditions of the Red-shafted Flicker (*Colaptes cafer*) species

**Species:** Cassin’s sparrow (*Peucaea cassinii*)

**Link:** https://ebird.org/species/casspa?siteLanguage=ht_HT

**Condition:** This bird was not seen anymore since 2017, perhaps because of climate change; emigrating before time to their locations for their summer stays.

**Species:** Rufous crowned sparrow (*Aimophila ruficeps*)

**Link:** https://ebird.org/species/rucspa

**Condition:** This species of sparrow prevails in the mountains. There is an adjacent mountain range called ‘Sierra de Orinda’, which may provide shelter for this bird. After 2013, it did not show up anymore.

**Species:** Black-chinned sparrow (*Spizella atrogularis*)

**Link:** https://ebird.org/species/bkcspa

**Condition:** It shown a similar behavior to the previous species, and not many individuals were observed from 2014 to 2017 (except for two). The monitoring routes studied correspond to their breeding areas. The presence of cattle in the monitoring routes may be an element of their behavior in the observations.

**Species:** Indigo bunting (*Passerina cyanea)*

**Link:** https://ebird.org/science/indbun

**Condition:** For some years, this species had a notable presence and similar to the previous species, its observations suddenly dropped.

**Species:** Common pauraque (*Nyctidromus albicollis*)

**Link:** https://ebird.org/species/compau

**Condition:** This bird is from coast areas. Hence, it is not common in this zone, but some individuals were observed, perhaps because they were moving between coasts. Similar to the species described above, they were not seen after 2013.

**Species:** *Aztec rail (*Rallus tenuirostris*)

**Link:** https://ebird.org/species/kinrai2?siteLanguage=es_MX

**Condition:** This species was not observed before 2009. It is in danger of extinction.

**Species:** Northern cardinal (*Cardinalis cardinalis)*

**Link:** https://ebird.org/species/norcar

**Condition:** The count of this species have significantly dropped, though at least some of them have been observed during the period. Nevertheless, their numbers have been decreasing, possibly because they have changed their nesting areas. It is a very desired bird by illegal traffickers.

**Species:** Chihuahua raven (*Corvus cryptoleucus*)

**Link:** https://www.audubon.org/field-guide/bird/chihuahuan-raven

**Condition:** This bird has reduced its numbers, especially during the last two years of monitoring. Perhaps they have changed their nesting locations.

**Species:** Red-shafted Flicker (*Colaptes cafer*)

**Link:** https://audubonportland.org/local-birding/kids-guide/backyard-birds/flicker

**Condition:** The behavior of this species may indicate seasonal patterns regarding changes in their nesting areas.

### 3.3. Species that increased in numbers

This section includes the analysis of three groups of species: pigeons, sparrows, hawfinch, and three isolated species, which had an increase in their numbers. In the case of the pigeons, four types of species were identified and their increase are summarized in Figure 16.

**Fig 16.**
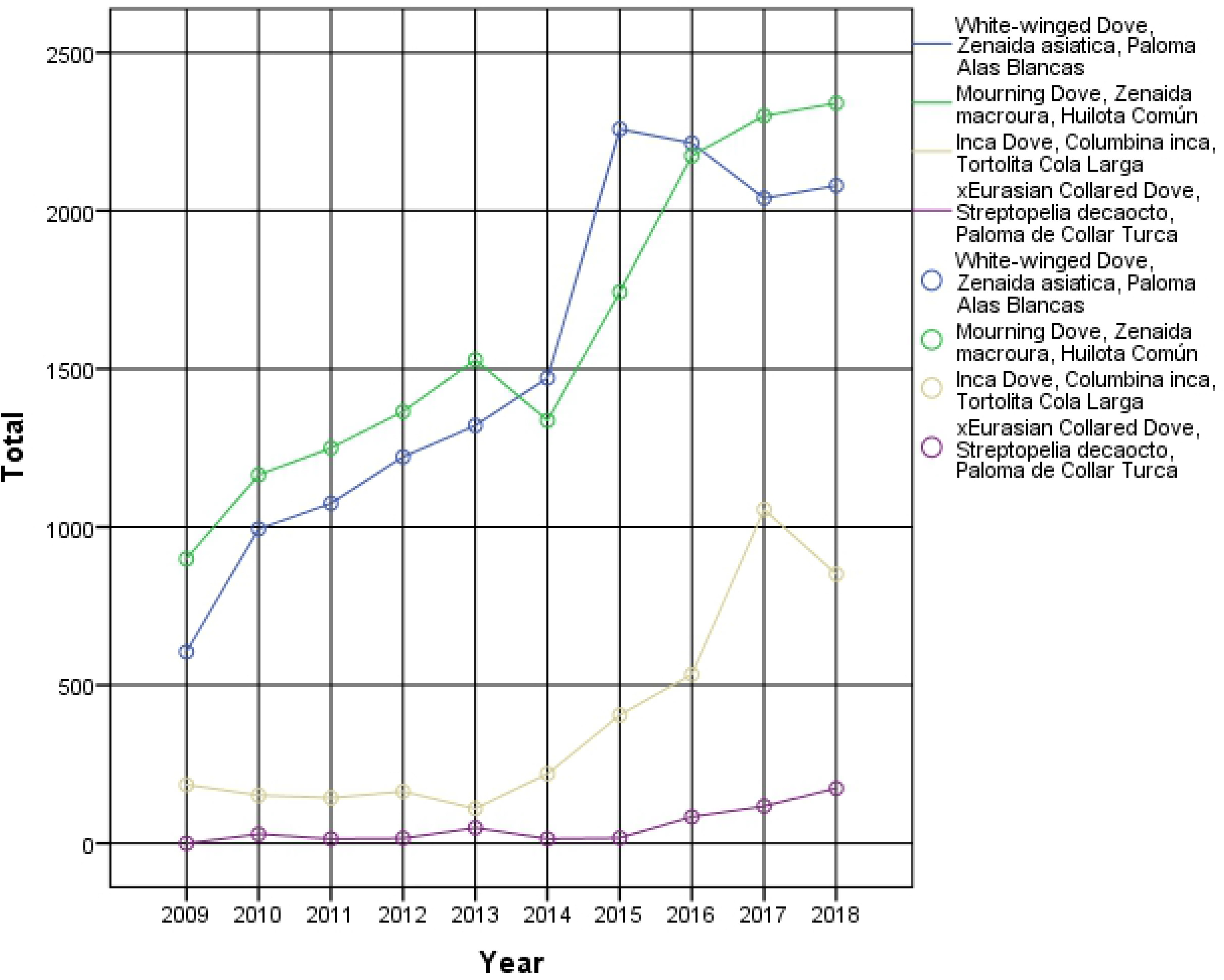
Increase of the four pigeon species.

**Species:** White winged dove (*Zenaida asiática*)

**Link:** https://ebird.org/species/whwdov?siteLanguage=es

**Species:** Mourning dove (*Zenaida macroura*)

**Link:** https://ebird.org/species/moudov

**Species:** Inca dove (*Columbina inca*)

**Link:** https://ebird.org/species/incdov

**Species:** Eurasian colored dove (*Streptopelia decaocto)*

**Link:** https://ebird.org/species/eucdov

According to the above results, it is not possible to clearly appreciate the increase of the *Eurasian Collared Dove species, as it is an invasive species. Therefore, it requires special attention, because the increase in their numbers produces a displacement of native species (see Figure 17).

**Fig 17.**
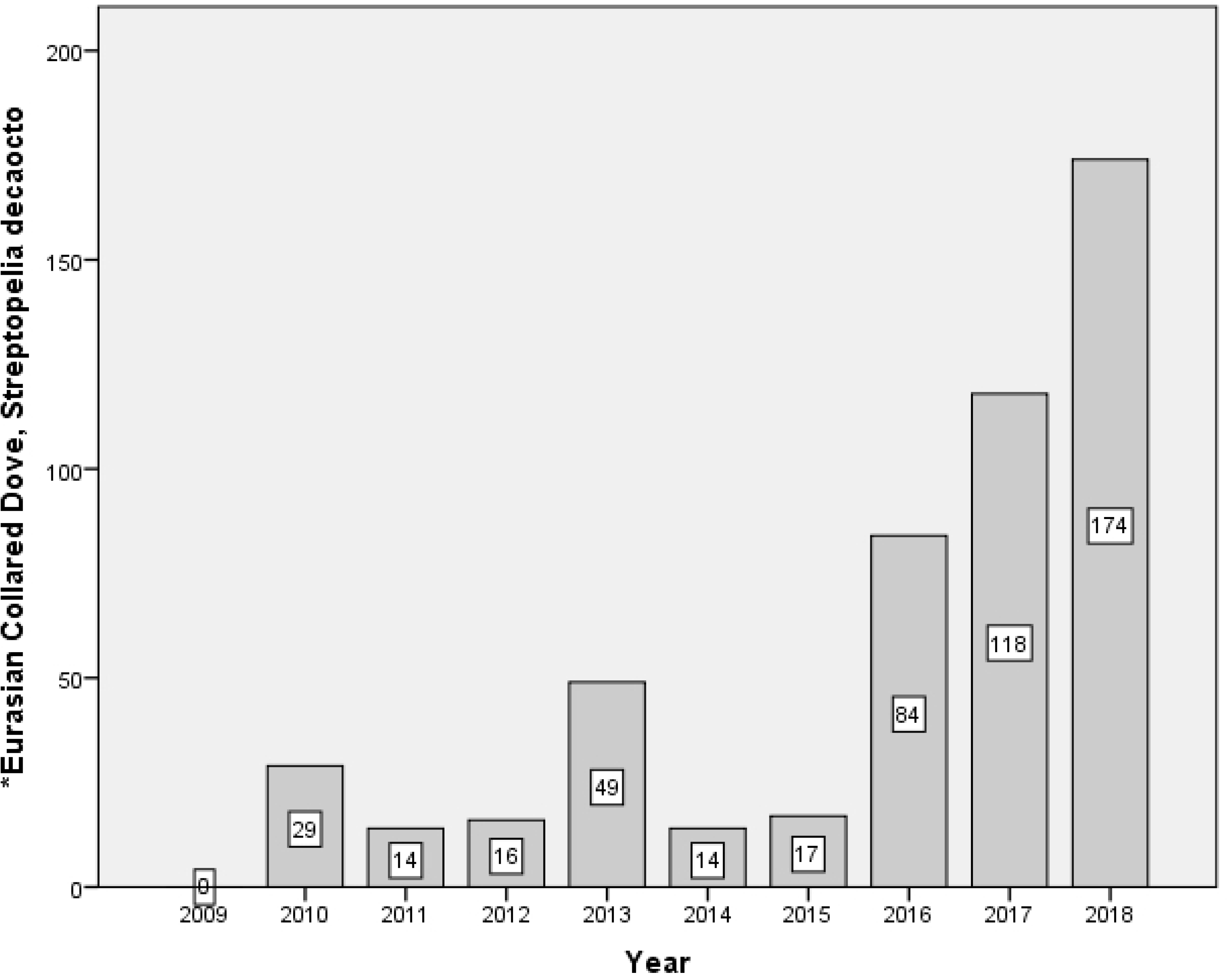
Increase of the *Eurasian Collared Dove (*Streptopelia decaocto*) species

According to the data gathered of the four pigeon species, a correlation analysis was carried out to identify significance and correlation values in all the cases, as Table 6 shows, where the correlations of these species indicate mutual protection, which means that their numbers increase together.

**Table 6.** Correlations of the four pigeon species that increased

|  |  | White winged dove | Mourning dove | Inca dove | *Eurasian collared dove |
| --- | --- | --- | --- | --- | --- |
| White winged dove | Pearson Correlation | 1 | .911** | .685** | .647** |
|  | Sig. (2-tailed) |  | .000 | .001 | .002 |
|  | N | 20 | 20 | 20 | 20 |
| Mourningdove | Pearson Correlation | .911** | 1 | .815** | .749** |
|  | Sig. (2-tailed) | .000 |  | .000 | .000 |
|  | N | 20 | 20 | 20 | 20 |
| Incadove | Pearson Correlation | .685** | .815** | 1 | .845** |
|  | Sig. (2-tailed) | .001 | .000 |  | .000 |
|  | N | 20 | 20 | 20 | 20 |
| Xeurasiancollareddove | Pearson Correlation | .647** | .749** | .845** | 1 |
|  | Sig. (2-tailed) | .002 | .000 | .000 |  |
|  | N | 20 | 20 | 20 | 20 |

\*\* Correlation is significant at the 0.01 level (2-tailed).

Regarding the sparrow species, they are coming closer and are identified more frequently in urban and semi urban areas, as they might be escaping from their natural predators, and they are feeding from the amount of waste produced in the cities, parks and gardens. The three species of sparrows that increased are (see Figure 18):

**Fig 18.**
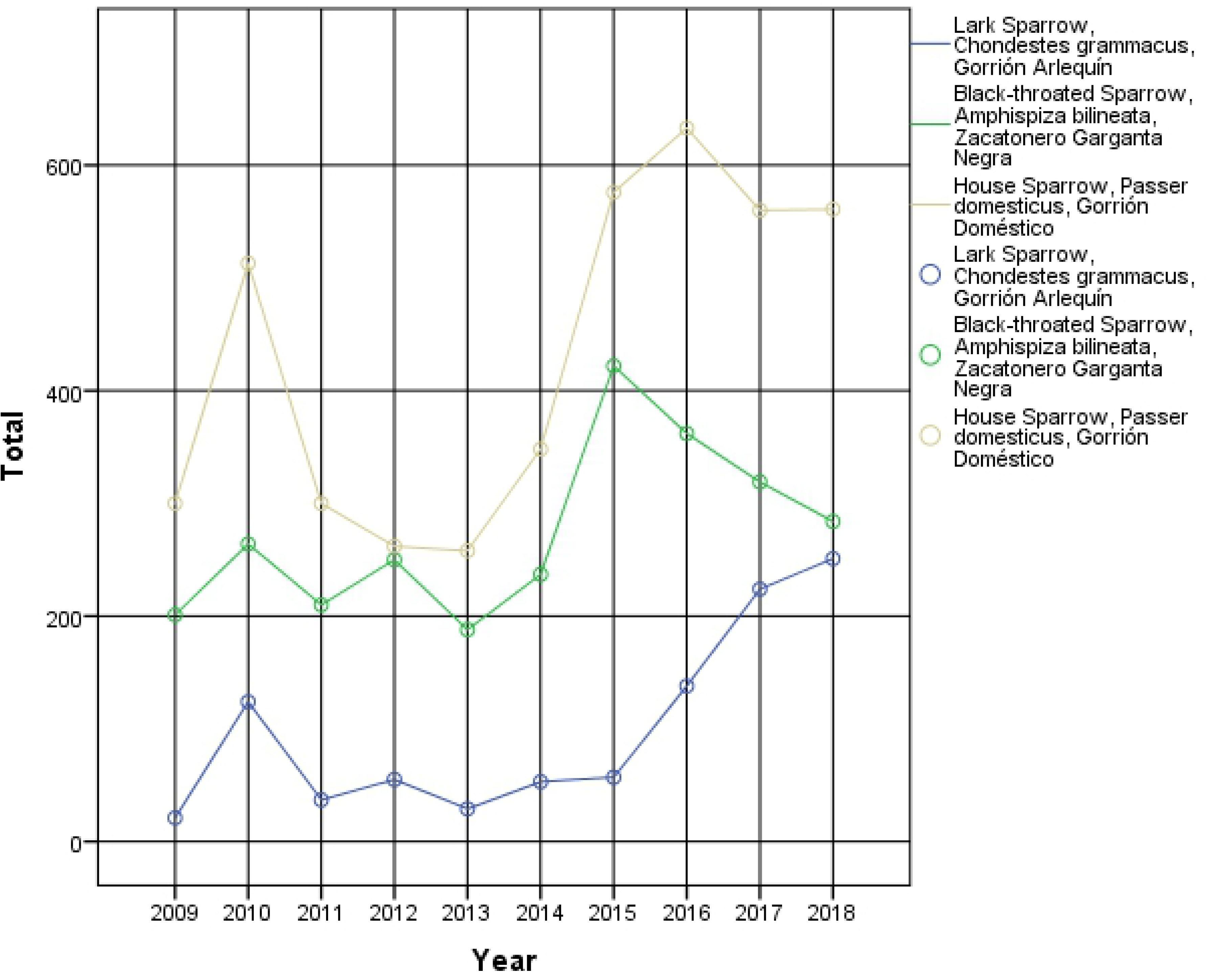
Increase of the three sparrow species

**Species:** Black throated sparrow (*Amphispiza bilineata*)

**Link:** https://ebird.org/species/bktspa

**Species:** Lark sparrow (*Chondestes grammacus*)

**Link:** https://ebird.org/species/larspa

**Species:** House sparrow (*Passer domesticus*)

**Link:** https://ebird.org/species/houspa

The observations of the two species of hawfinch that increased were characterized by similar foraging habits and distributions. These are associated to areas with urban trees, such as the mulberry tree. Figure 19 shows the two hawfinch species that increased their numbers.

**Fig 19.**
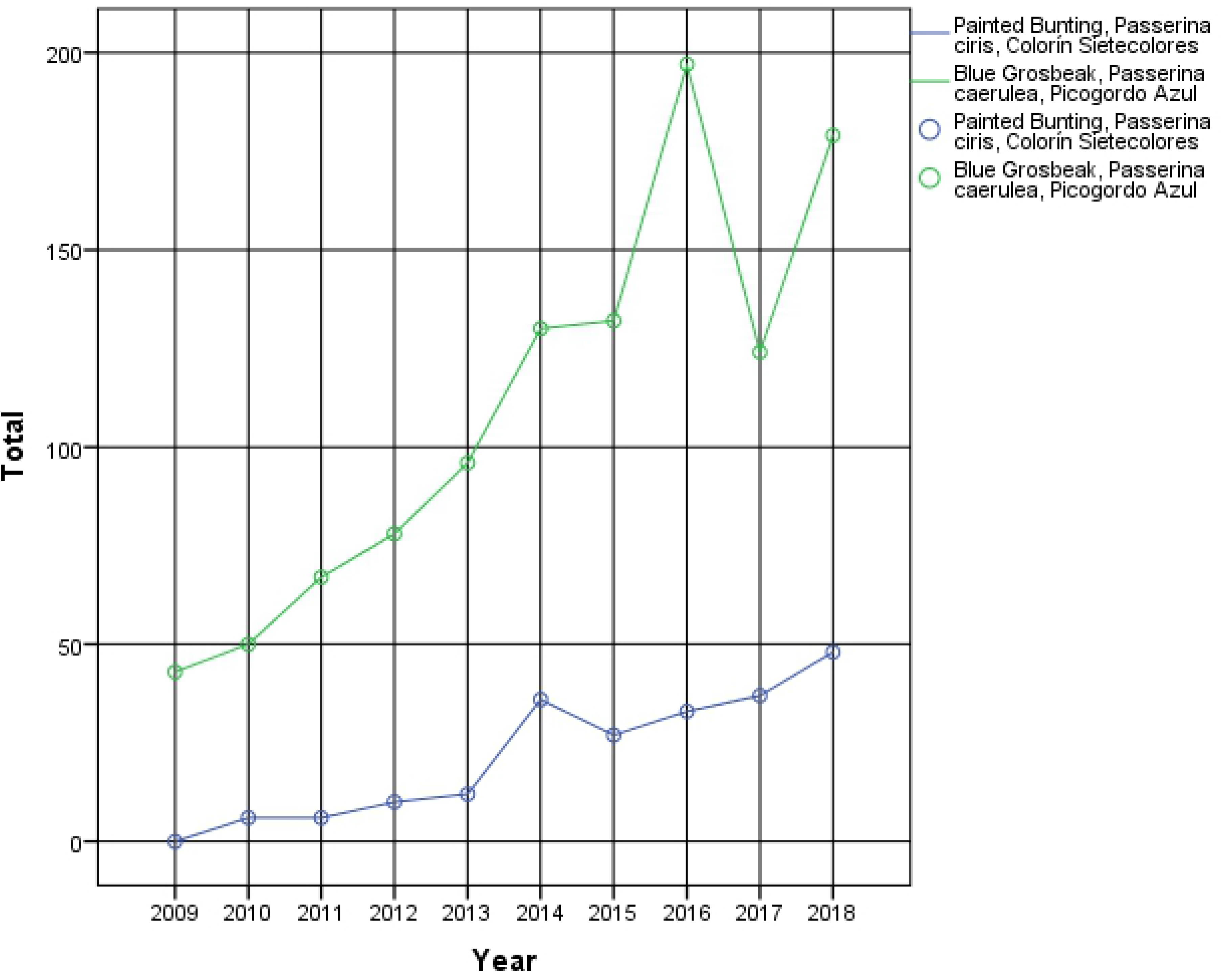
Increase of the two hawfinch species

**Species:** Blue grosbeak (*Passerina caerulea*)

**Link:** https://ebird.org/species/blugrb1

**Species:** Painted bunting (*Passerina ciris*)

**Link:** https://ebird.org/species/paibun

The results observed in the three species are shown below and separately by species.

**Species:** Red Winged Blackbird (*Agelaius phoeniceus*)

**Link:** https://ebird.org/species/rewbla

**Condition:** This species is associated to agricultural activities. Hence, its continuous increase in the analyzed routes is expected, as Figure 20 shows.

**Fig 20.**
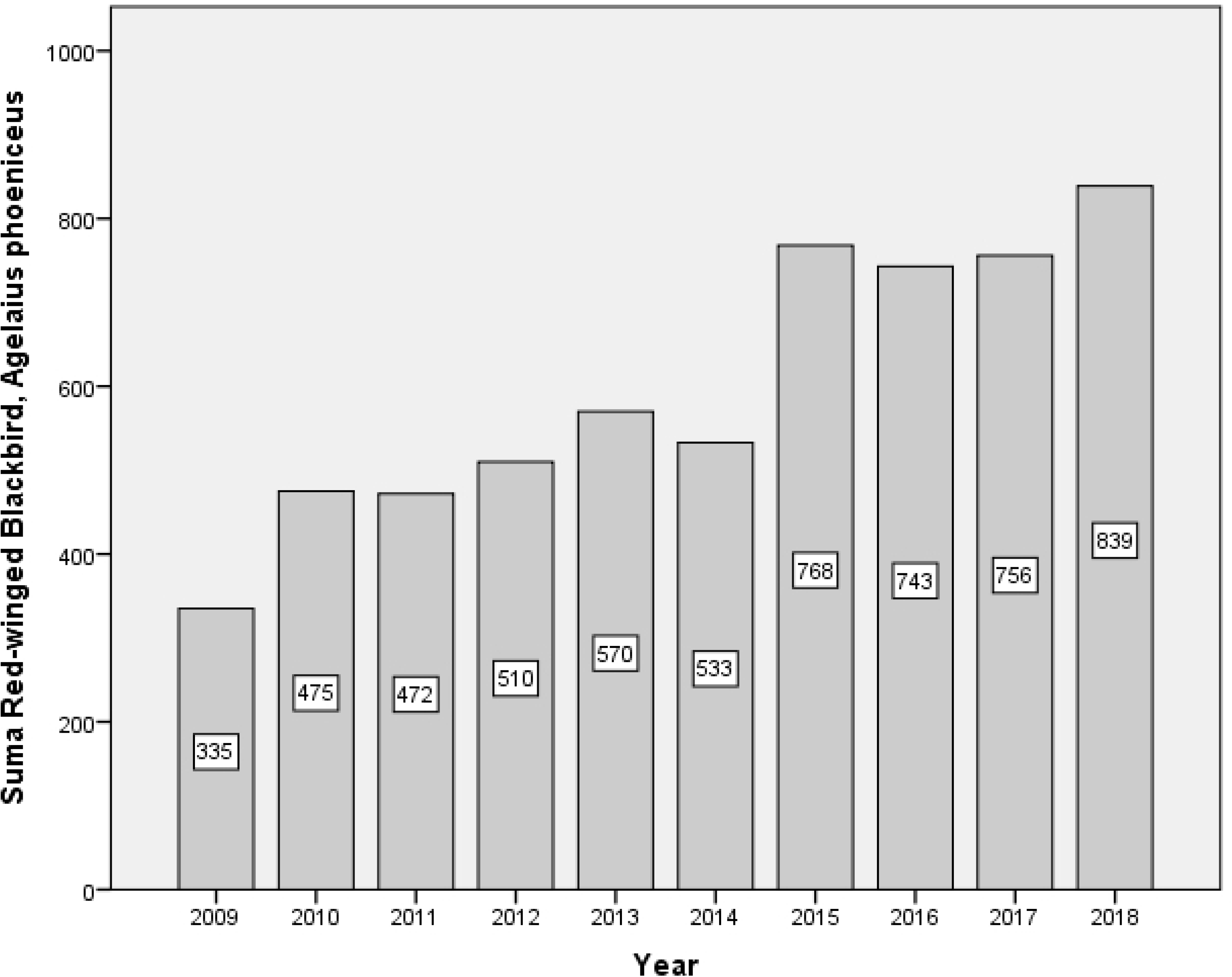
Increase of the Red winged blackbird (*Agelaius phoeniceus*) species

**Species:** Vermilion flycatcher (*Pyrocephalus rubinus*)

**Link:** https://ebird.org/species/verfly

**Condition:** This species has currently increased, probably due to the presence of insects coming from the agricultural activities within the monitoring areas (see Figure 21).

**Fig 21.**
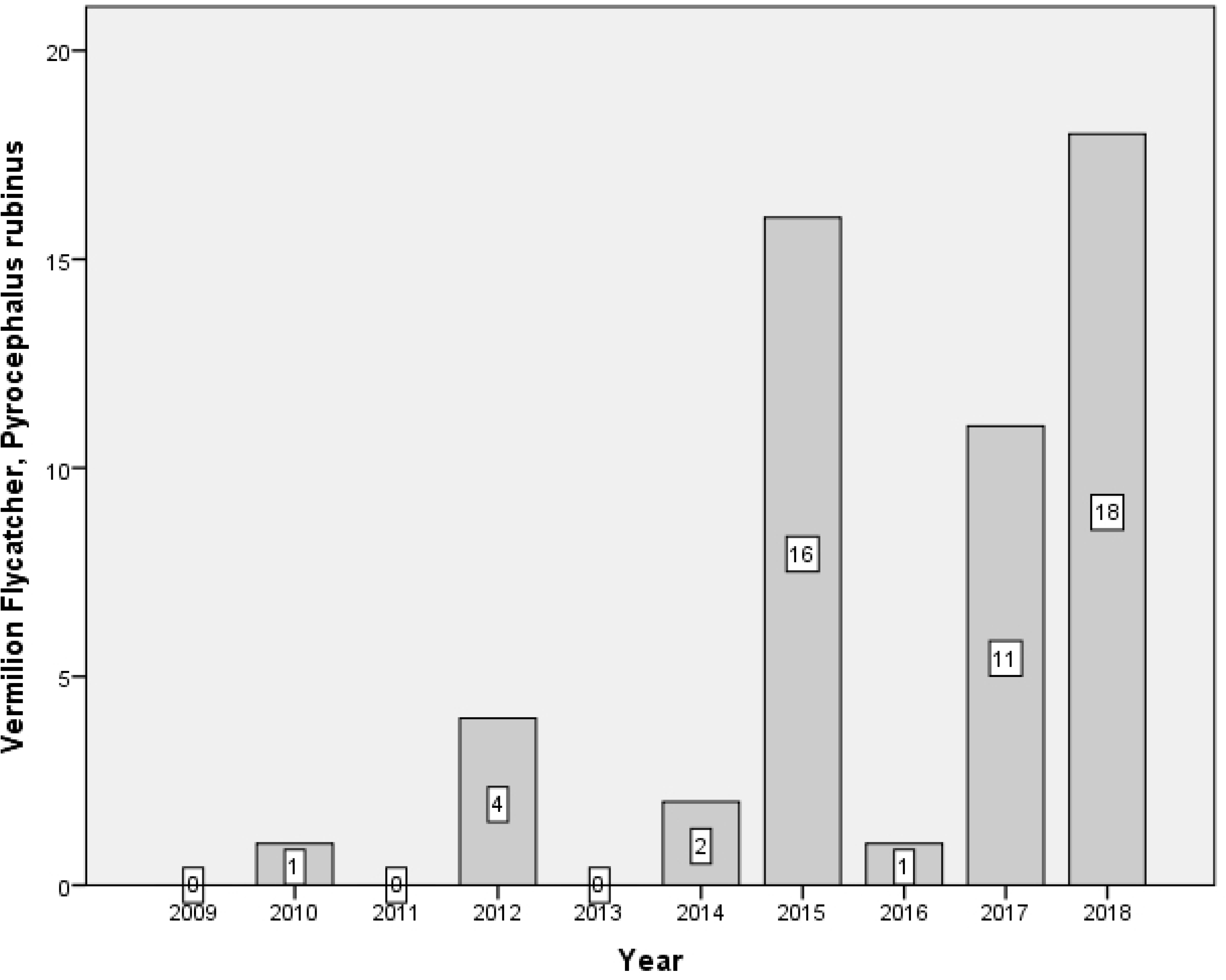
Increase of the Vermilion flycatcher (*Pyrocephalus rubinus*) species

**Species:** *Verdin (*Auriparus flaviceps*)

**Link:** https://ebird.org/species/verdin?siteLanguage=es_MX

**Condition:** This species is regaining its population, probably thanks to the reforestation of native plant species in the monitoring areas (see Figure 22).

**Fig 22:**
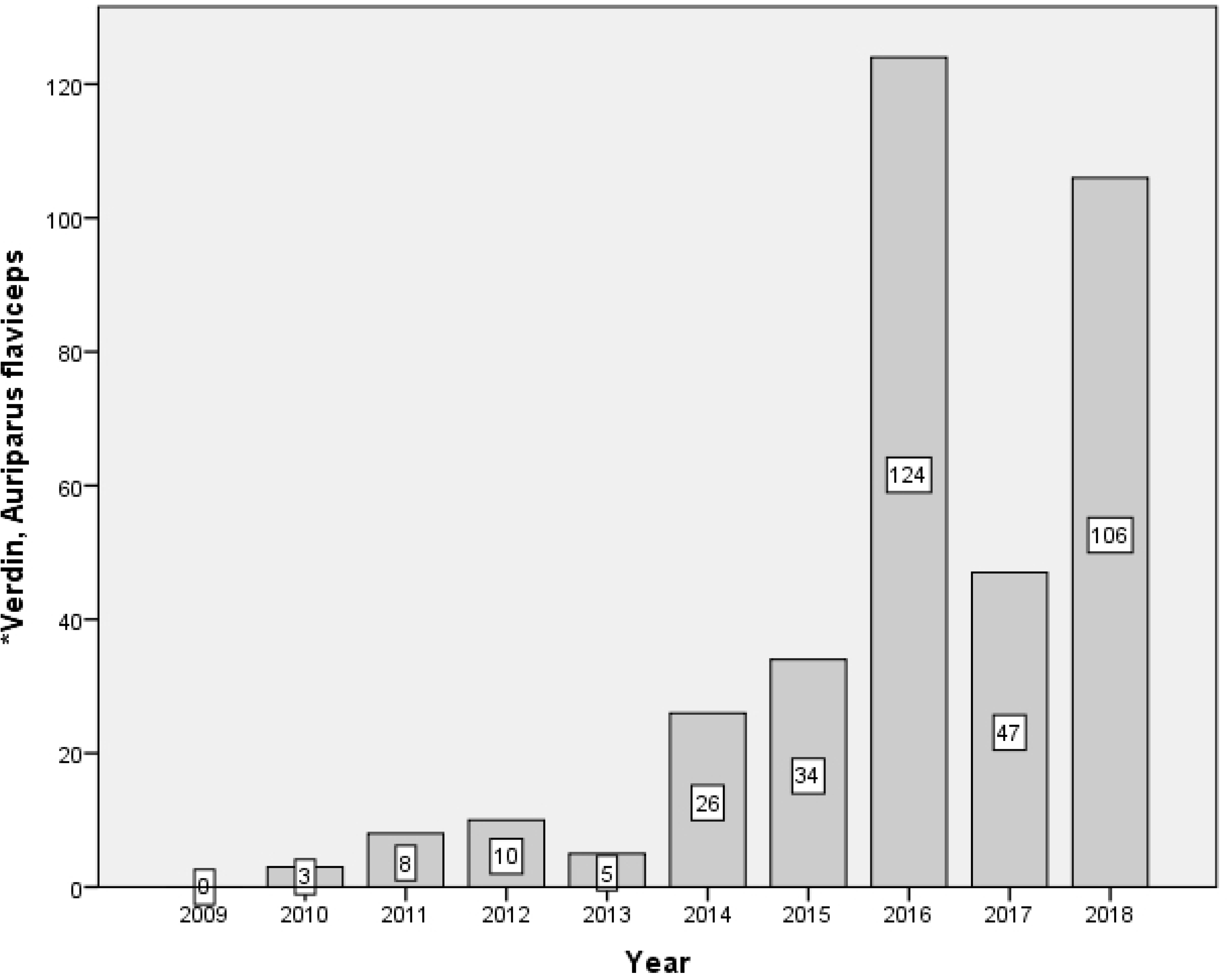
Increase of the *Verdin (*Auriparus flaviceps*) species

### 3.4. Analysis by number of species

In this study, the total number of species observed, during the evaluated period of 2009-2018 and in both routes, showed a global decrease from 99 to 76 species (23.33%). Figure 23 shows the yearly decrease of species for both routes.

**Fig 23.**
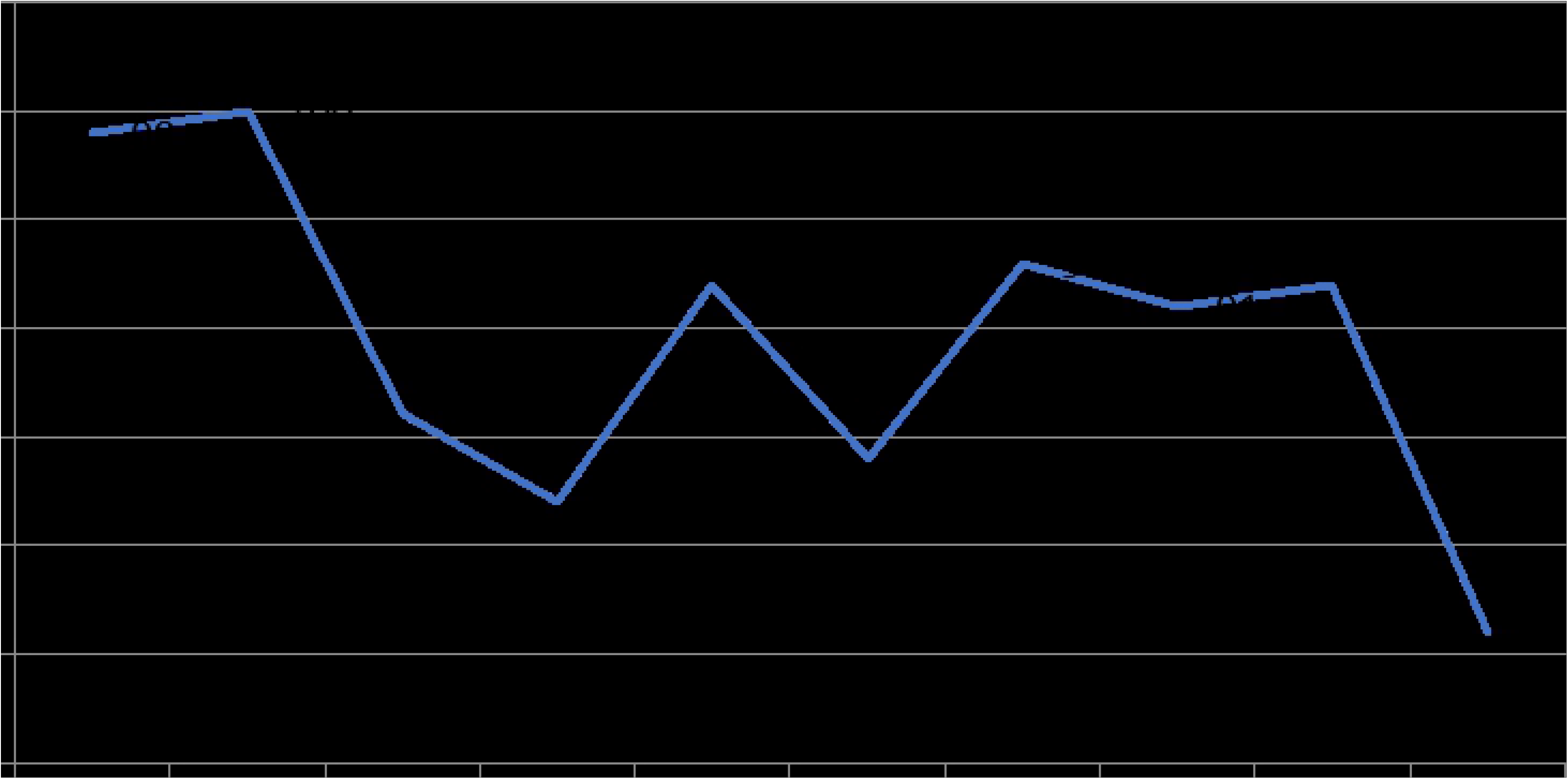
Decrease in the total number of species in both routes.

During 2009, the first year of monitoring, 76 different species were observed in Santa Monica route, and there was a progressive reduction until 58 species were observed during 2018 (23.6% decrease). In La Regina route, the observations during the same period of time resulted in a reduction from 91 to 70 species, which represented an almost identical reduction in percentage: 23.07% (see Figure 24).

**Fig 24.**
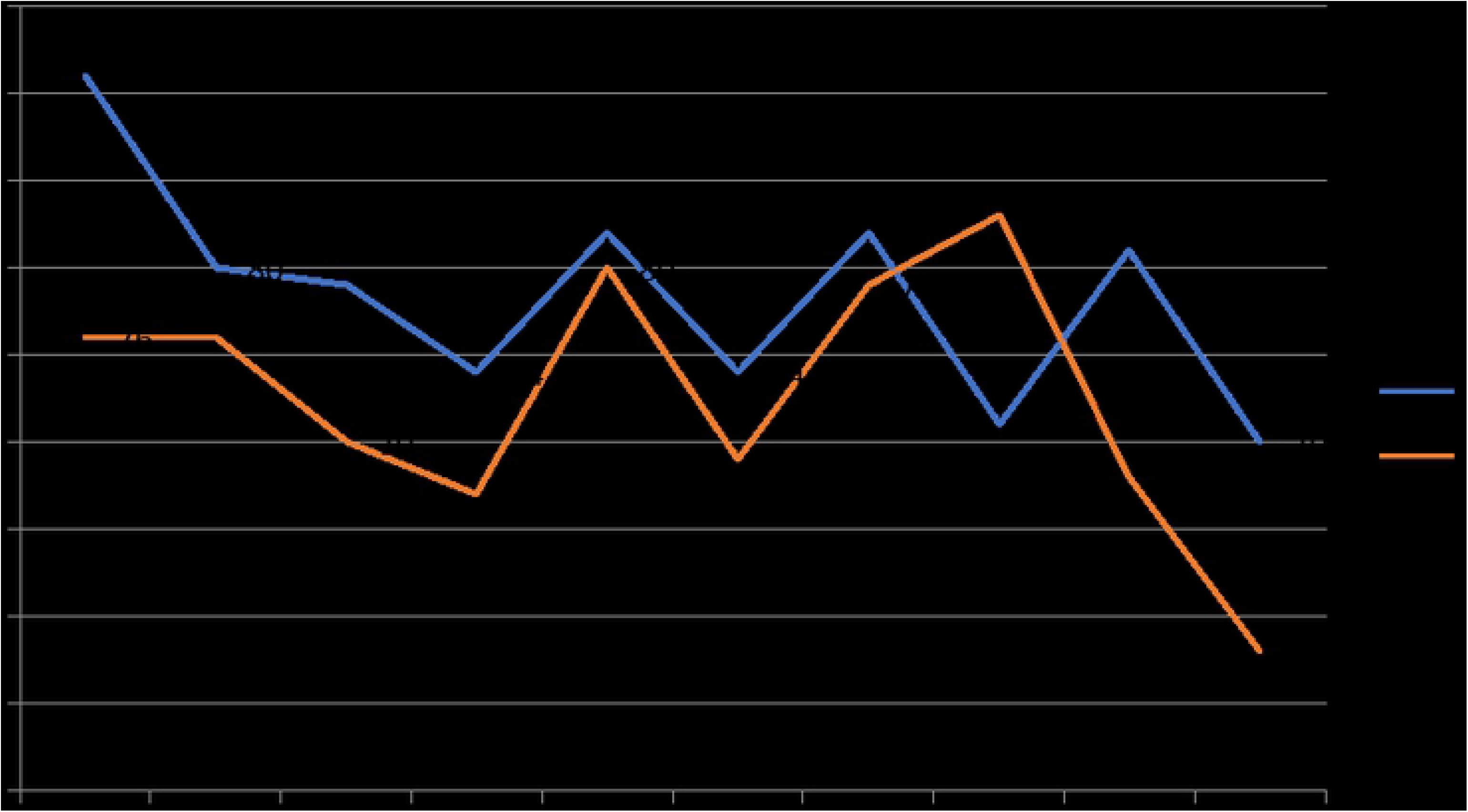
Comparison of species’ decrease by route

### 3.5. Other findings from observations

The results included in this section are relevant for monitoring, because they may be useful for birdwatchers, in the sense that they communicate the ideal conditions in which the studied species can be found in the region. Such conditions were drawn from correlations between species and climatological conditions (see Table 7).

**Table 7.** Correlations between species and climatological conditions

| Species | Climatological conditions |
| --- | --- |
| Says poheby | It is observed more easily with low temperatures and cloudy skies. |
| Ash-throat flycatcher | It is observed in the morning, with more frequency when the temperature is high and the wind is low. |
| Couch's kingbird | Positive correlation with the windy conditions of the sky. |
| Blue grosbeak | Direct positive correlation with temperature. |
| Western meadowlark | Negative correlation between temperature at the end of the route and positive correlation with the rate of windy sky. |
| White-tailed kite | Negative correlation between sightings and wind speed. |
| Elf owl | Very positive correlation between windy conditions and sightings. |
| Blue-winged teal | Very high positive correlation between windy conditions and sightings. |
| Yellow-rumped warbler | Very high positive correlation between windy conditions and sightings. |

## 4. Discussion and conclusions

According to the results, new species in the area were identified in the monitoring (Rufous-crowned sparrow, Ruddy-ground dove and Common-ground dove), which are out of their migratory routes; a condition that are attributed to changes in their usual routes and of the climatological conditions, which force them to seek alternative routes. Different species observed were only present during some years and their presence was not registered again. Changes in their migration routes puts some species in several risks, both at rural as in urban environments, mainly due to the presence of different predators.

The Aztec rail species, of which some sightings were reported since 2009, drew the attention of national and international ornithologists, because they were previously observed at the center of Mexico (very far from Chihuahua). This Mexican species is in danger of extinction, according to the Mexican Official Standard 059 from the Secretariat of the Environment and Natural Resources [39].

The frequency of sightings has increased for some species thanks to the insects linked with agricultural activities within the area. Invasive species like the Eurasian collared-dove and the European starling have notoriously increased their populations, which cause the displacement of native species.

Some positive activities have been accomplished in the monitoring routes, such as the reintegration of native species. This has turned out to be beneficial to some species, such as the Verdin, which is regaining its population. In fact, a change in the counting of different species between the years 2013-2014 was observed. The species whose numbers dropped or that disappeared during the monitoring period have similar habits related with meadows or lower lands. Conversely, the species that have increased their numbers seem to have the tendency to fly or remain in higher areas.

We can pose the following hypotheses regarding the decrease in numbers: the indiscriminate use of agrochemicals without performing environmental evaluations may negatively affect bird numbers; this has been documented in news outlets [40]; and the area is notably studied because of reports about contamination with high levels of arsenic and fluorine [41]. This information has been constantly checked by the local media for being connected to high rates of cancer in the inhabitants of the region [42].

The data and information provided by birdwatching which supported this research is important for decision-making regarding environmental issues, although there is no evidence of such use at a local, regional or even national level. There is a need for spreading and communicating results of the kind discussed in this article. We argue that such a need is imperative, due to the variations in the sightings of species, which evidence the decrease and increase of certain species and may lead to environmental concerns; these are issues that should be part of an educational program for everyone within the region [43].

Formal education about environmental issues provides the opportunity to connect the traditional knowledge of communities (holders of a profound knowledge about environmental surroundings) and scientific knowledge, for which universities may act as intermediaries. Activities related to the environment and the results provided by this research may help to expand the global-local roles in the ecosystem processes of nature and species’ migratory movements, as well as alleviating the human and socio-environmental issues that currently affect us.

The existence of dissemination processes for such kinds of results and the creation of formal and informal educational programs are important to raise awareness about the environment and work toward improving it and making it more sustainable. Moreover, we propose that we need to go beyond, as two fundamental necessities have to be addressed: a) the creation of scientific observatories, as publicly accessible systems to retrieve environmental and ecological data and information; and, as a result b) defining mechanisms for promoting the generation of citizen science, with the participation of both scientists and the civil society.

The extensive data used in this research, which is already recorded, can be considered as a starting point to implement a scientific observatory about birdwatching through the appropriate and clear presentation of the data and deriving useful information from it, which will allow people and institutions to take decisions based on the search and discrimination of the relevance of such data [44] in an ecological information system. The proposal of such system has the aim to continue with its compilation, treatment and diffusion through the use information technologies, whose effects can be demonstrated in the reflection and knowledge about the topic and for aiding environmentally friendly decision-making, which avoids putting in danger the ecosystems. Such kinds of initiatives require the participation of formal institutions like governments, universities or scientific centers for their organization.

Birdwatching activities in Mexico, as in the state of Chihuahua, offer enough conditions to those people involved for, almost in an empiric way, constituting an important example in the development of citizen science, for gathering data, and disseminating knowledge related to these issues. In this context, citizen science has been increasingly used to collect biodiversity data and to inform the management and preservation of the environment [45].

In other contexts, the concept of citizen science has been used as a way to democratize scientific knowledge [46]. Moreover, it represents important actions that can be undertaken, especially when financing turns out to be limited, irregular or inexistent; therefore, it becomes a reliable and feasible alternative for monitoring species [47].

Citizen science as an alternative: (i) encourages a scientific culture and brings science to the society; (ii) expects to make citizen scientists, through the participation of volunteers who gather or process data for research and decision-making; and (iii) provides more value to the citizen observation capacity than that of any sophisticated equipment by itself [48]. The citizen, without being a trained scientist, becomes a prosumer (producer-consumer of information).

## 5. Acknowledgements

The authors wish to thank the CONABIO, the United States Geological Survey Patuxent Wildlife Research Center and the Canadian Wildlife Service Research Centre for inviting citizens to be part of this research, and for providing the necessary training and the technical facilities to conduct this important volunteer activity.

